# How Binding Site Flexibility Promotes RNA Scanning in TbRGG2 RRM: A Molecular Dynamics Simulation Study

**DOI:** 10.1101/2024.09.25.614920

**Authors:** Toon Lemmens, Jiri Sponer, Miroslav Krepl

## Abstract

RNA Recognition Motifs (RRMs) are a key class of proteins that primarily bind single-stranded RNAs. In this study, we use unbiased molecular dynamics simulations to obtain insights into the intricate binding dynamics between uridine-rich RNAs and TbRGG2 RRM. Complementing structural experiments that unveil a primary binding mode with a single uridine bound, our simulations uncover two supplementary binding modes where adjacent nucleotides encroach upon the binding pocket. This leads to a unique molecular mechanism through which TbRGG2 RRM is capable of rapidly transitioning the U-rich sequence. In contrast, presence of non-native cytidines induces stalling and destabilization of the complex. By leveraging extensive equilibrium dynamics and large variety of binding states, TbRGG2 RRM effectively expedites diffusion along the RNA substrate while ensuring robust selectivity for U-rich sequences despite featuring a solitary binding pocket. Using recently developed Stafix potential, we substantiate our description of the complex dynamics by simulating fully spontaneous association process of U-rich sequences to the TbRGG2 RRM. Our study highlights critical role of dynamics and auxiliary binding states in interface dynamics employed by RNA-binding proteins, which is not readily apparent in traditional structural studies, but could represent a general type of binding strategy employed by many RNA-binding proteins.

## Introduction

The RNA recognition motif (RRM) is a common RNA-binding protein domain in eukaryotes, capable of recognizing various RNA sequences with high fidelity (1). This makes the RRM-containing proteins a ubiquitous feature of RNA metabolism, including expression, splicing, editing, and stabilization (2). Up to 1% of human genes include RRMs (3) and their widespread utilization in eukaryotic evolution stands in stark contrast to their relatively simple and conserved topology. Namely, a single RRM domain is typically composed of ∼90 amino-acids with a consensual secondary structure of β_1_α_1_β_2_β_3_α_2_β_4_ (4), where two α-helices fold against one side of the antiparallel β-sheet surface. While its structure shows limited variations among the different proteins, the RRM’s specificity for different RNA sequences can be astoundingly tuned via substitutions of its surface-exposed amino acids (4-11). This ability is further enhanced by the diverse modes of protein-RNA recognition the RRMs can facilitate. Namely, the exposed β-sheet surface of RRM is the most common binding site for RNAs (12), but in principle any part of the domain can be tuned for RNA recognition, including the α-helices, the loops and the terminal chains (9,10,13-18). Usually two to five nucleotides are directly recognized by a single RRM (3). The RNA-binding proteins can contain multiple RRM domains or act on a single substrate as multimers, further increasing the specificity of the RRM-RNA recognition. The widespread role of RRM domains in RNA metabolism made them a staple of protein-RNA structural biology and numerous structures of RRM protein-RNA complexes were determined by X-ray crystallography and NMR spectroscopy (2). A limitation of these studies is that they basically provide a static ensemble-averaged picture of the bound protein-RNA complex in equilibrium. In reality, the RRM-RNA interaction, even at equilibrium, would be more accurately described as a dynamic ensemble of conformations sampled with different lifetimes and populations (11,19-22). Likewise, the structural details of the binding process (binding pathways) cannot be inferred from structural studies of fully bound complexes (23).

Molecular dynamics (MD) simulations are a computational method for studying movement of biomolecules using carefully calibrated set of empirical potentials commonly known as the force fields (*ff*s). Explicit-solvent standard MD can be used to routinely observe thermal fluctuations of the entire protein-RNA complex on a microsecond-scale with essentially infinite spatial and temporal resolution (24-26). This level of detail is currently inaccessible for experimental methods and the simulations thus fill an important gap in our knowledge of the biomolecular systems. However, the MD is limited by the affordable length of the calculation and adequacy of the utilized *ff*s (24). The latter can be particularly challenging for description of the protein-RNA complexes as the *ff*s are generally optimized for simulations of isolated proteins and nucleic acids. One such *ff* issue is overcompaction of single-stranded RNA molecules (ssRNA) with the standard RNA *ff*s (27-29), as they tend to overwhelm the intermolecular interactions and disrupt protein-ssRNA complexes (30,31). We have recently tackled this problem by developing a *ff* modification called Stafix which eliminates the overcompaction and significantly improves the *ff* performance in protein-ssRNA simulations. Stafix even allows comprehensive studies of the binding process of simple protein-ssRNA complexes starting from the fully unbound state (23).

The TbRGG2 is a protein responsible for kinetoplastid RNA (kRNA) editing in the kinetoplastid protists, such as the *Trypanosoma brucei* (32). During the kRNA editing, specific uridine insertions or deletions are performed on the mitochondrial mRNAs, the extent of which varies among the transcripts, ranging between few to hundreds of uridines (33,34). The TbRGG2 alters the mRNA structure, a process which was shown as essential for successful kRNA editing (35). The TbRGG2 is composed of two domains, the N-terminal glycine-rich region and C-terminal RRM (36); the latter is studied in this work. The TbRGG2 RRM stands out among the other RRMs in several ways. Firstly, it exhibits a rare multimodal binding by specifically recognizing both the U-rich and G-rich RNAs via separate binding sites. Secondly, the binding site for the U-rich sequences consists of a single binding pocket recognizing one uridine (Figure 1, A). The binding site of the G-rich sequences is currently unknown (37). Note that TbRGG2 is sometimes referred to as RESC13 in the literature (38,39).

**Figure 1:**
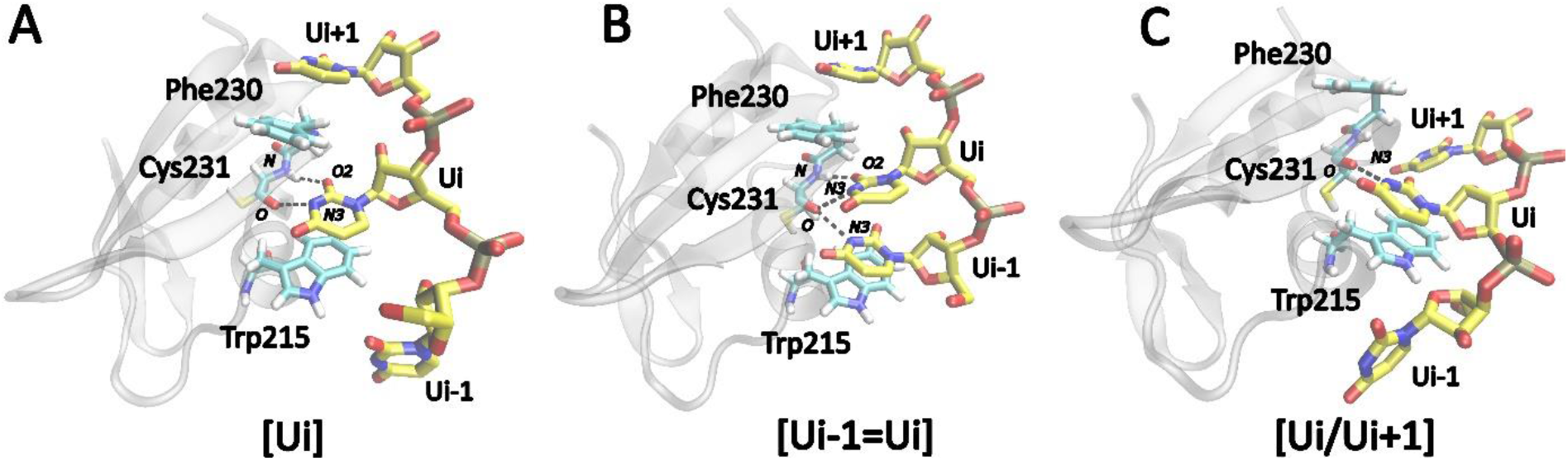
Representative structures of the three dominant states of TbRGG2 RRM-poly(U) binding interface as observed in MD simulations. The H-bonds characterizing each state are indicated with black dashed lines. The protein secondary structure is shown as transparent gray ribbon and the RNA and binding pocket amino acids are colored yellow and cyan, respectively. (A) Single uridine (observed also in the experiment), (B) the vertical and (C) horizontal states. To annotate the three states, we use symbolic representations of binding pocket structure, where “[“ and “]” refer to the clamp formed by the Trp215 and Phe230 side-chains, respectively, and “=“ and “/” indicate vertical and horizontal arrangements of the uridines, respectively. The “i-1”, “i+1”, and “i±1” indices indicate upstream, downstream, and either of the nucleotides, respectively. This annotation is utilized throughout the study. Where relevant for the description, “i” is replaced with a specific uridine residue number.

Here we have used MD simulations to study the μs-scale equilibrium dynamics of the complex between TbRGG2 RRM and poly-(U) ssRNA. We show that the dynamics of the binding site is characterized by large-scale conformational fluctuations including fully reversible loss of the specific interactions. The key observation is that the main binding mode (shown by experiments) can be transiently replaced with two alternative binding modes in which two uridines simultaneously occupy the pocket. Strikingly, the newly observed binding modes are highly selective for uridines as well. Such structural dynamics of the interface allows smooth sliding of the RNA along the protein and we sometimes observed complete spontaneous shift of the poly-(U) sequences all the way to the strand termini. An encounter with non-native cytidines tends to cause either swift transition back to U-rich regions or disengagement of the RNA from the binding pocket, followed by another binding attempt elsewhere on the sequence. We also carried out spontaneous binding simulations starting from the unbound state, where we observed successful formation of the native binding state as indicated by the experiments, followed by the characteristic equilibrium dynamics. Ultimately, these simulations revealed the same behavior as simulations initiated from the bound state, confirming the observed equilibrium dynamics is quite converged and not biased by the starting structure. We suggest that dynamic binding mechanisms such as the one employed by TbRGG2 RRM could constitute so far underappreciated element of protein-RNA recognition, largely invisible to conventional structural experiments due to its transient nature (40), but with potentially significant importance and utility in tuning protein-RNA interactions.

## Methods

### Selection and preparation of initial structures

We have used the X-ray structure of the T. brucei TbRGG2 RRM domain complexed with the 5′-UUU-3′ (3U) RNA (chain B/K; PDB: 6E4P) as the starting structure, with the individual uridines numbered from U1 to U3 (37). An additional RRM domain (Chain A) was used in a limited amount of simulations (see Supplementary Information). In some simulations, we have extended the RNA chain at both ends using Pymol to obtain complexes with bound 5′-UUUUU-3′ (5U) RNA, adding uridines U0 and U4 at the 5′- and 3′-ends, respectively. As starting structures for some of the 5U complex simulations, we also used simulation snapshots where two uracils simultaneously occupy the binding pocket. By replacing individual nucleotides with cytidines in the 5U complex, we obtained starting structures with bound RNA sequences of 5′-UCUUU-3′, 5′-UUUCU-3′, 5′-UUCUU-3′, 5′-CCUCC-3′, or 5′-CCCCC-3′. In some simulations, TbRGG2 RRM was modified by replacing the K219 and K232 residues with alanines. Simulations of the free protein (chain B; PDB: 6E4P) were performed by removing the RNA from the complex structure.

The initial structures for the spontaneous binding simulations from the unbound state (23) were prepared by placing the TbRGG2 RRM protein and the 5′-UUUU-3′ RNA ∼20 Å apart. This distance was shown to be sufficient for randomizing the ssRNA’s internal structure and the first spontaneous point of contact between the two biopolymers (23). For the spontaneous binding simulations, the initial RNA structure was prepared with Nucleic Acid Builder and corresponded to an A-RNA helix with the complementary strand removed. Poly-(U) sequence of four nucleotides was selected to increase sampling efficiency compared to longer sequences and to reduce off-pathway binding (23). The uridines were in this case counted from U1 to U4. Lastly, initial coordinates for the simulations of the putative RRM homodimer with and without RNA bound were obtained from chains A/B/K and A/B (or A/D) of the 6E4P structure, respectively (Supplementary information).

### System building, force fields and simulation protocol

We utilized the tLeap program of AMBER 20 (41) to generate the initial simulation files. The RNA was described with the OL3 *ff* (42) and we additionally applied the recently developed Stafix potential (23), which eliminates spurious RNA self-interactions via extensive rescaling of the intra-RNA van der Waals interactions (see also Supplementary Information). Stafix greatly improves accuracy of the simulations of bound protein-ssRNA complexes and is indispensable for simulations of spontaneous binding of ssRNA to proteins from the unbound state (23). The applied Stafix factor for all bound simulations and most of the spontaneous binding simulations was 0.5. A factor of 0.1 was also tested for some of the spontaneous binding simulations. The RRM was described by the ff12SB *ff* (43); note that for RRM protein-RNA complexes we prefer to use the ff12SB version over the ff14SB for reasons explained elsewhere (23). The biomolecules were immersed in an octahedral box of SPC/E water molecules (44) with minimal distance of 12 Å between the solute and the edge of the simulation box. Some of the binding simulations were also performed with the OPC water model (45). Physiological excess-salt ion concentration of 0.15 M was obtained by adding KCl ions (46) at random positions around the solute. The equilibration of the systems was performed using the standard protocol (47) in the pmemd.MPI program of AMBER 20. Afterwards, we ran production simulations of varying length (2–10 μs) using the GPU-enabled pmemd.cuda program (48). In all cases, multiple parallel simulations of each system were carried out. SHAKE (49) and hydrogen mass repartitioning (50) were applied in all simulations, allowing a 4 fs integration step. Long-range electrostatics was handled by the particle mesh Ewald (51) in a periodic boundary conditions setup. The distance cut-off for calculation of the non-bonded Lennard-Jones (LJ) interactions was 9 Å. Langevin thermostat and Monte Carlo barostat (41) were used to regulate the production simulations.

### Analyses

We have used cpptraj (52) and VMD (53) to analyze and visualize the trajectories, respectively. Graphs and molecular figures were prepared with Gnuplot and Inkscape, respectively. To monitor the state of the TbRGG2 RRM binding pocket, we visually inspected every trajectory. The state of the pocket was manually annotated based on the number of occupying nucleotides, their mutual orientation and identity. The resulting data is presented as horizontal bar graphs with the individual states indicated by different colors. Apart from visual analysis, we also separately evaluated the state of the pocket using an automated method where the distinction is made based on the H-bonds formed between the nucleotides and the Cys231(O) and Cys231(N) atoms, with cut-off distance of 3.5 Å between donors and acceptors and acceptor-hydrogen-donor angle of 135°. A second criterium was the distance between Trp215(CE3) and Phe230(CG) atoms, with distances below and above 9.0 Å corresponding to the single state and the vertical/horizontal states, respectively. The automated method was used to quickly evaluate the state of the pocket in each simulation, however, due to enormous complexity of the binding pocket dynamics, we ultimately used manual annotation to produce the graphs below.

## Results

Below, we present results of multiple-microsecond explicit-solvent MD simulations of the TbRGG2–poly(U) complex. We also explore complexes involving RNA sequences with varying content of cytidines or mutated protein domains. Finally, we present the results of spontaneous binding simulations where we have studied the entire association process of the TbRGG2-poly(U) complex, starting from the unbound state. The number of conducted simulations is 60 with a cumulative length of 524 μs. The results below describe the key observations derived from our analyses of the simulated complexes. Simulations of isolated RRM and its putative homodimers are described in Supplementary Information.

### Temporary accommodation of two uridines at the binding site is the hallmark of the TbRGG2 RRM complex dynamics

The experimental X-ray structure (PDB: 6E4P) shows one uridine stacked between the binding pocket residues Trp215 and Phe230 (Figure 1A), with the base specifically recognized by U2(N3)-Cys231(O) and Cys231(N)-U2(O2) H-bonds (35). We call this arrangement ***single uridine*** state. All the MD simulations initiated from this structure revealed significant (sub)μs-scale dynamics involving conformational transitions and temporary interruptions of the binding. Importantly, we have regularly observed two additional binding states in which two uridines simultaneously occupy the pocket. Based on the mutual orientation of the two bases within the pocket, we defined them as the ***vertical*** and ***horizontal*** state. The vertical state is formed by two uridines sandwiched between Trp215 and Phe230, giving rise to a quadruple stack (Figure 1B). Note that it may involve both upstream and downstream uridine with respect to the originally bound single uridine. In the horizontal state, the first uridine is sandwiched between Trp215 and Phe230, while the second (downstream) uridine is interacting with a hydrophobic patch on the RRM’s surface. This places both bases in roughly the same plane (Figure 1C And Supplementary Figure S1). Instances in which non-consecutive uridines formed either the horizontal or vertical states also appeared, albeit rarely. The hallmark of dynamics of the TbRGG2–poly(U) complex on microsecond timescale are frequent exchanges between the single uridine state, and the vertical and horizontal states, sometimes temporarily interrupted by an ***unbound*** state where no uridines are present within the binding pocket even though the RNA remains attached to the protein surface in proximity of the binding pocket.

### TbRGG2 specifically recognizes uridines in all three binding states

The single uridine state involves the Cys231(O)–Ui(N3) and Cys231(N)–Ui(O2) H-bonds. These interactions also align the uridine base for optimal stacking with the Trp215 and Phe230 aromatic side chains (Figure 1A). Such arrangement is not sterically feasible for larger purine bases while cytidine lacks compatible H-bond donors and acceptors. Strikingly, the specificity for uridines is maintained also for the vertical and horizontal binding states observed in our simulations. Namely, in the vertical state, the second uridine Ui-1 is also engaged in the Cys231(O)–Ui-1(N3) H-bond, resulting in bifurcated H-bonding (Figure 1B). In the horizontal state, the uridine Ui located deeper in the binding pocket is recognized similarly to the single uridine state while the additional downstream uridine Ui+1 is recognized by Val228(O)–Ui+1(N3) H-bond. There is also a fluctuating H-bond between Lys219(NZ) and Ui+1(O4) (Supplementary Information). In the vertical state, the distance between the aromatic rings of Trp215 and Phe230 increases by ∼1.0 Å on average compared to the single uridine state, reflecting the accommodation of the additional nucleotide. The distance similarly increases by ∼0.5 Å for the horizontal state. For details of the H-bond populations in all three states see Supplementary Information.

### Accommodation of two uridines by TbRGG2 RRM facilitates fast diffusion of the bound RNA

The experimental structure of the complex has only three nucleotides (U1-U3) resolved, with the middle U2 bound in the pocket. In our simulations of this structure, we observed either the U1 or U3 terminal nucleotides eventually entering the pocket as well, forming the vertical or horizontal states (Figure 1). In some cases, the U2 would later depart the pocket, leaving the U1 or U3 as the single nucleotide bound. This corresponds to sliding of TbRGG2 RRM along the bound RNA by a single step, raising a possibility of an intriguing stepwise nucleotide shift mechanism with the vertical and horizontal states acting as intermediates. We next performed simulations with extended 5′-UUUUU-3′-RNA bound (see Methods), to explore this potential shift mechanism further. In these simulations, we regularly observed multiple such transitions in either direction on the simulation timescale, supporting the existence of this spontaneous diffusion mechanism (Figure 2, Table 2). In some cases, we were able to observe shift of the entire bound RNA sequence all the way to the U0 or U4 terminal nucleotides (Supplementary Information). In conclusion, our simulations strongly suggest that TbRGG2 RRM is capable of rapid linear diffusion along the bound poly-(U) RNA by utilizing the vertical and horizontal binding states as intermediates. A shift by a single nucleotide can be completed as quickly as 0.5 μs, counting from the formation of the intermediate. As no biasing force was applied to facilitate the transitions, they randomly proceeded in upstream or downstream directions. The diffusion was further sped up by temporary disruptions of the binding, occasionally leading to no nucleotide being present in the binding pocket (the unbound state). This sometimes allowed the protein to “jump over and skip” some nucleotides by reattaching further up-or downstream.

**Figure 2:**
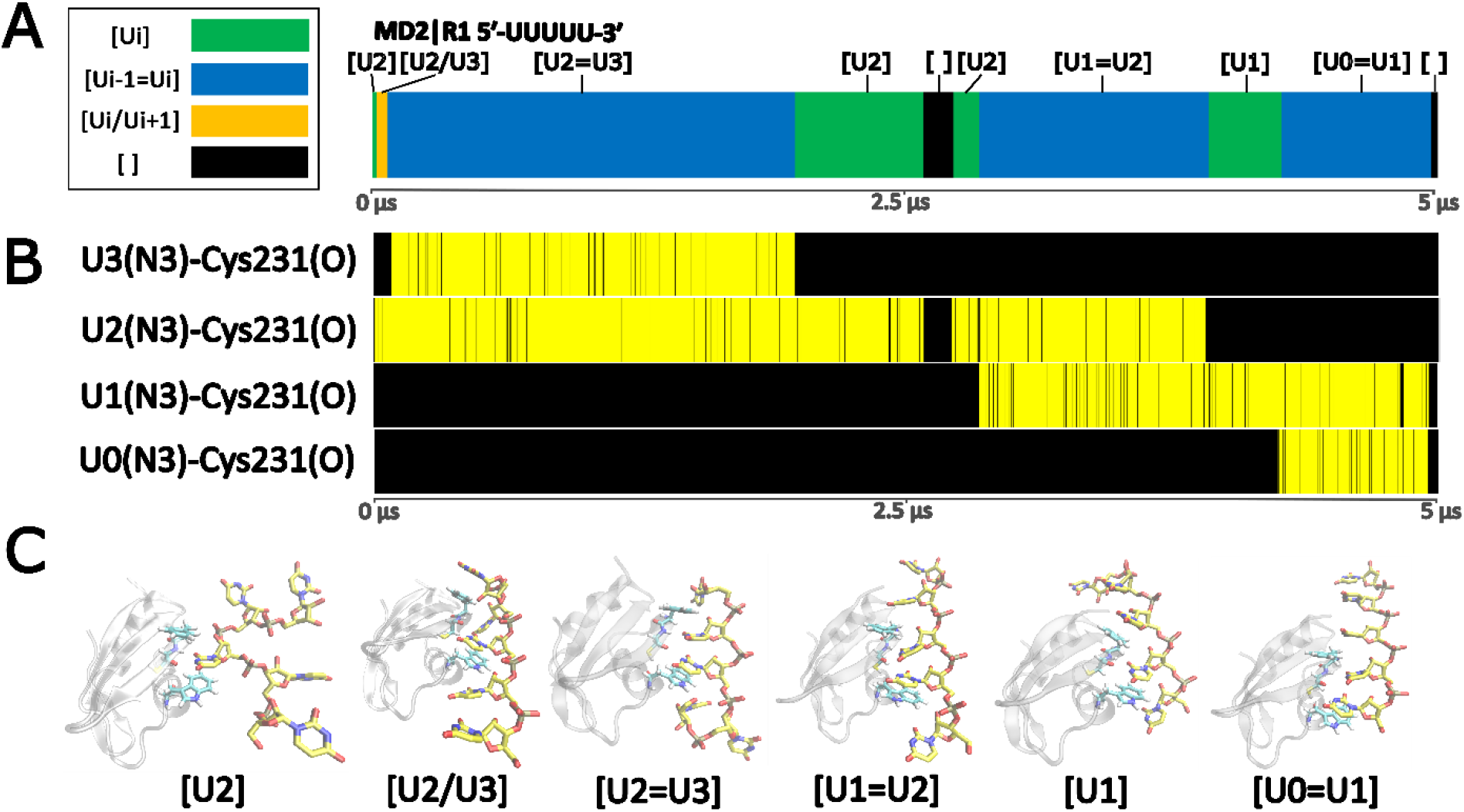
Example of TbRGG2 RRM diffusion along the 5′-UUUUU-3′ RNA. A time-development analysis of the MD2|R1 trajectory (see Supplementary Information for the description of all the other trajectories). (A) Graph of the binding pocket occupancy. The different binding states are color-coded according to the legend on the left. “[ ]” indicates no uridine is present in the binding pocket. (B) Time-development of the characteristic H-bond formed between the N3 atoms of the occupying uridines and Cys231(O). Yellow and black indicate presence and absence of the H-bond, respectively. (C) Snapshots from the simulation showing representative structures for each of the observed states.

**Table 1.**
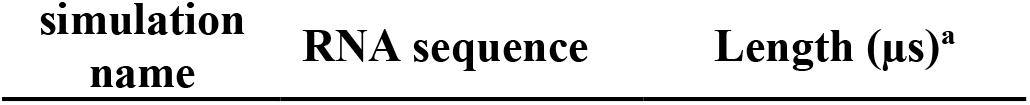
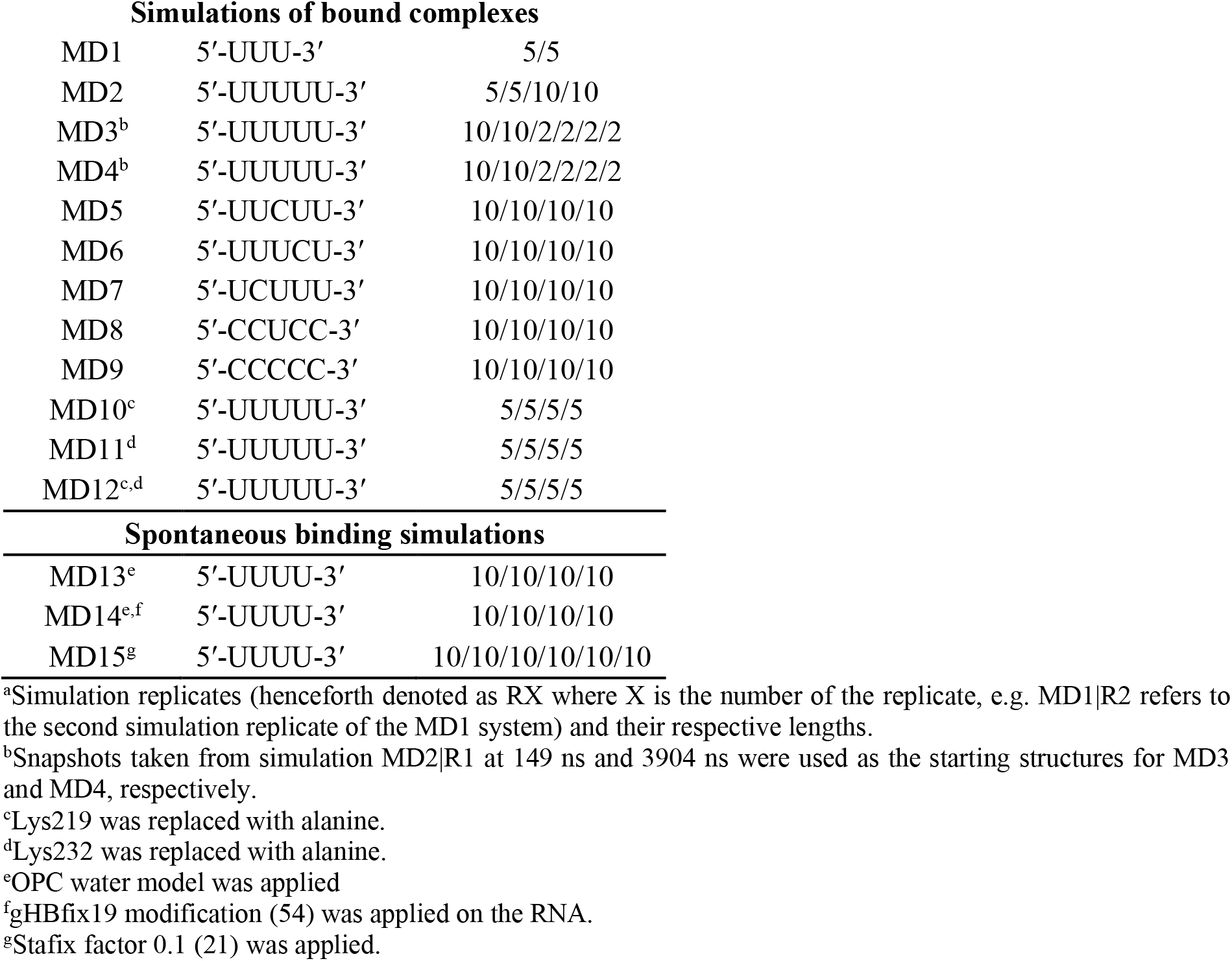
List of simulations.

**Table 2.** Number of transitions observed in MD simulations of the TbRGG2–5’-UUUUU-3’ complex.

| Simulation | Starting state<sup>a</sup> | 5'<sup>b</sup> | 3'<sup>b</sup> | Simulation length [μs]<sup>c</sup> |
| --- | --- | --- | --- | --- |
| **Simulations of bound complexes** |  |  |  |  |
| MD2\|R1 | [U2] | 4 | 1 | 5 |
| MD2\|R2 | [U2] | 2 | 2 | 5 |
| MD2\|R3 | [U2] | 3 | 1 | 10 |
| MD2\|R4 | [U2] | 0 | 1 | 10 |
| MD3\|R1 | [U2=U3] | 7 | 7 | 10 |
| MD3\|R2 | [U2=U3] | 3 | 3 | 10 |
| MD3\|R3 | [U2=U3] | 3 | 3 | 2 |
| MD3\|R4 | [U2=U3] | 1 | 1 | 2 |
| MD3\|R5 | [U2=U3] | 0 | 0 | 2 |
| MD3\|R6 | [U2=U3] | 0 | 0 | 2 |
| MD4\|R1 | [U1=U2] | 6 | 5 | 10 |
| MD4\|R2 | [U1=U2] | 1 | 3 | 10 |
| MD4\|R3 | [U1=U2] | 1 | 1 | 2 |
| MD4\|R4 | [U1=U2] | 2 | 0 | 2 |
| MD4\|R5 | [U1=U2] | 3 | 3 | 2 |
| MD4\|R6 | [U1=U2] | 1 | 0 | 2 |
| **Spontaneous binding simulations** |  |  |  |  |
| MD13\|R1 | - | 3 | 4 | 10 (0.35) |
| MD13\|R2 | - | 5 | 4 | 10 (0.13) |
| MD13\|R3 | - | 9 | 8 | 10 (0.95) |
| MD13\|R4 | - | 4 | 3 | 10 (0.18) |
| MD14\|R1 | - | 1 | 1 | 10 (0.81) |
| MD14\|R2 | - | 7 | 9 | 10 (0.44) |
| MD14\|R3 | - | 8 | 8 | 10 (0.41) |
| MD14\|R4 | - | 0 | 0 | 10 (-) |
| MD15\|R1 | - | 0 | 0 | 10 (-) |
| MD15\|R2 | - | 0 | 0 | 10 (0.05) |
| MD15\|R3 | - | 5 | 5 | 10 (0.02) |
| MD15\|R4 | - | 0 | 0 | 10 (1.20) |
| MD15\|R5 | - | 4 | 3 | 10 (5.59) |
| MD15\|R6 | - | 0 | 0 | 10 (-) |

<sup>a</sup>Type of binding pocket interaction in the initial structure of the simulation. Simulations MD13-MD15 were the spontaneous binding simulations, with the two molecules completely separated at the start.
<sup>b</sup>Number of transitions in upstream or downstream (5' and 3') directions; it refers to any change in the identity or number of uridines occupying the binding pocket. For the spontaneous binding simulations, this was counted once the first successful binding attempt was observed.
<sup>c</sup>For spontaneous binding simulations, the number in parentheses refers to the time of the first successful binding. When no number is stated, no successful binding was observed on the simulation timescale.

### Cytidine tends to be rejected by TbRGG2 RRM during the diffusion process

Above, we have described the mechanism which allows TbRGG2 RRM to quickly diffuse along the poly-(U) RNA (Figure 2). To determine how the RRM would respond to an encounter with a non-uridine nucleotide, we explored multiple systems where selected uridines of the bound 5′-UUUUU-3′ RNA were mutated to cytidines (see Methods). Firstly, we explored the 5′-UUCUU-3′ system, making the C2 the nucleotide bound in the pocket at the simulation start. This mutation abolishes the critical Cys231(O)–Ui(N3) H-bond, causing an immediate misalignment of the cytidine with respect to the Trp215 and Phe230 side chains (Figure 3). Soon after the start of the simulation, one of the neighboring uridines entered the pocket, forming a state best described as a *hybrid state*, a combination of the vertical and horizontal states (Figure 3 and Supplementary Information). In all 5′-UUCUU-3′ simulations, the initially bound cytidine was ultimately irreversibly expelled from the binding pocket and supplanted by neighboring uridines (Supplementary Information).

**Figure 3:**
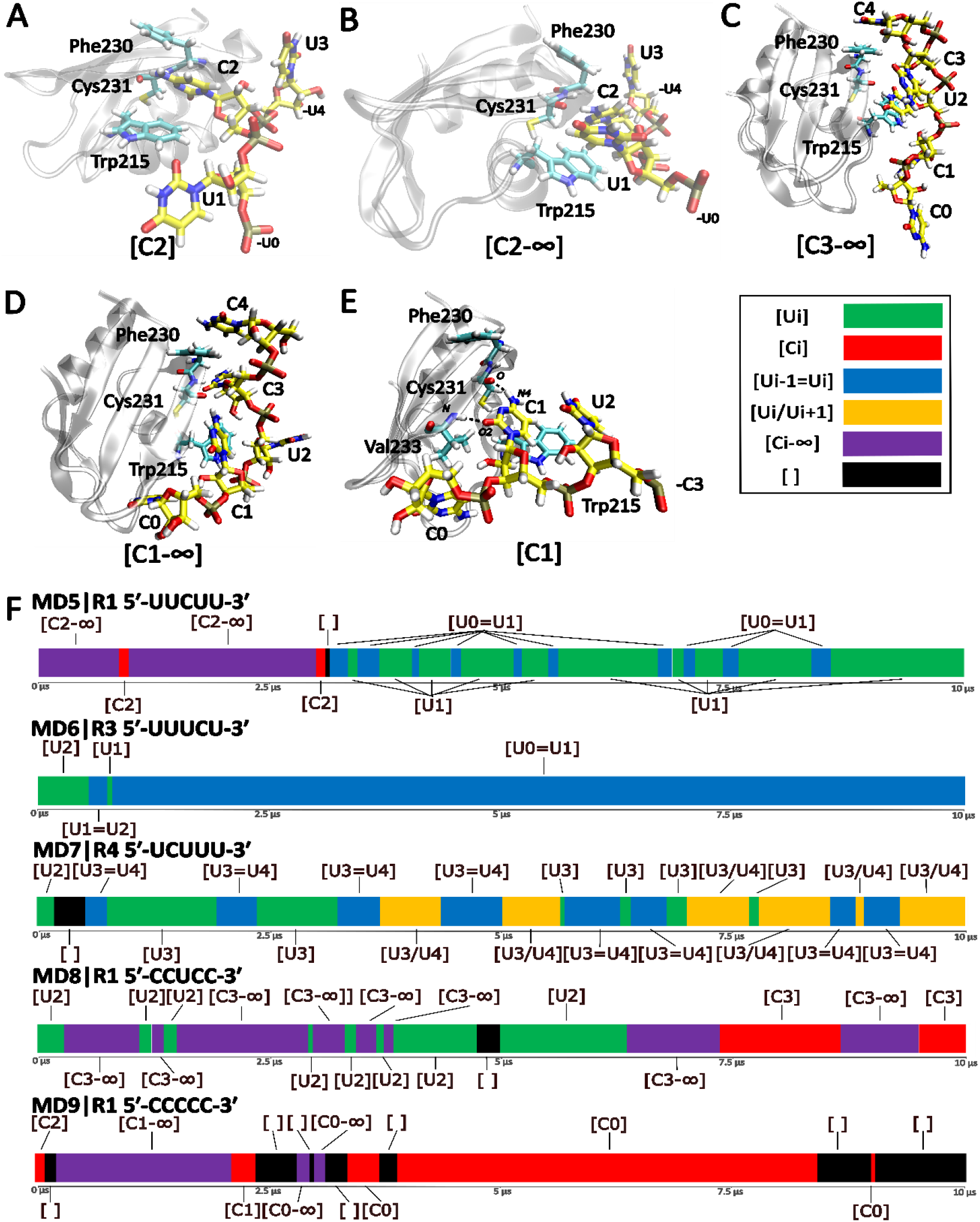
Simulations of TbRGG2 RRM bound to cytidine-containing RNAs. (A) The starting structure of MD5 simulations with 5′-UUCUU-3′. (B, C) Two variants of the hybrid state, where uridine and cytidine simultaneously occupy the pocket. (D) A state with only the backbone of the cytidine entering the binding pocket. (E) One of the arrangements attempted by the system to form an interaction with the cytidine. Due to enormous structural richness of the binding states involving cytidine, we collectively label all of them as “Ci-∞”, referring to a variety of states in which the protein tries to achieve stable binding of the cytidine by simultaneously forming interactions with uridine, another cytidine or part of the RNA-backbone. Binding of a single cytidine is labelled as Ci. (F) Time-development of the binding pocket occupancy in the individual cytidine-containing systems (MD5 to MD9). A single representative simulation of each system is shown (see the Supplementary Information for the rest).

Secondly, we explored the effect of placing cytidine immediately upstream or downstream of the single bound uridine. We subsequently observed attempted shifts in both directions, but those towards the cytidine almost universally failed, resulting ultimately in the RNA shifting towards the uridine-rich portion of the RNA. In other words, in the 5′-UCUUU-3′ systems, the RNA tended to shift towards the 3′-end while the opposite occurred for the 5′-UUUCU-3′ simulations. In one case, we observed the RNA separating completely from the protein during an attempted shift in the direction of the cytidine, followed by rebinding further along the RNA chain, thus entirely skipping the non-native cytidine. Thirdly, we tried mutating all the uridines except U2 to cytidines, i.e. the 5′-CCUCC-3′ sequence, thus potentially making the shifts unfavorable in both directions. Indeed, we observed the U2 located in the binding pocket being regularly invaded by the flanking C1 or C3 (Figure 5B And 5C). However, attempts to occupy the binding site by a cytidine were unstable, followed either by quick restoration of the sole U2 binding or by dissociation of the RNA from the binding pocket. For the sake of completeness, we also explored behavior of the 5′-CCCCC-3′ system. In this case, the system initially tried to explore numerous conformational arrangements involving the cytidines, however, none of them was stable and the RNA eventually left the binding pocket entirely.

### Specific protein mutations hamper the diffusion mechanism

In our simulations, we observed the side-chain of Lys232 influencing the dynamics and orientation of the uridines occupying the binding pocket, by H-bonding with the Ui(O4) atoms (Figure 4A). Similarly, the Lys219 side-chain located on the opposite side of the binding pocket from Lys232 often formed H-bonds with the Ui+1 nucleotide of the horizontal state (Figure 4B). To determine how the Lys232 and Lys219 residues impact the characteristic dynamics of the TbRGG2 RRM binding pocket and the diffusion scanning mechanism, we prepared complexes where they were replaced by alanines. The Lys232Ala mutation led to increased flexibility of the nucleotide in the binding pocket and the diffusion mechanism was essentially abolished, with only a single greatly unstable vertical state observed (Supplementary Information). The uridines in the binding pocket could still be replaced by adjacent uridines, but these replacements occurred solely via temporary dissociation of the RNA from the binding pocket followed by a new binding. In conclusion, the simulations suggest the Lys232 residue is important for properly aligning the nucleotides in the binding site as well as enabling the diffusion mechanism.

**Figure 4:**
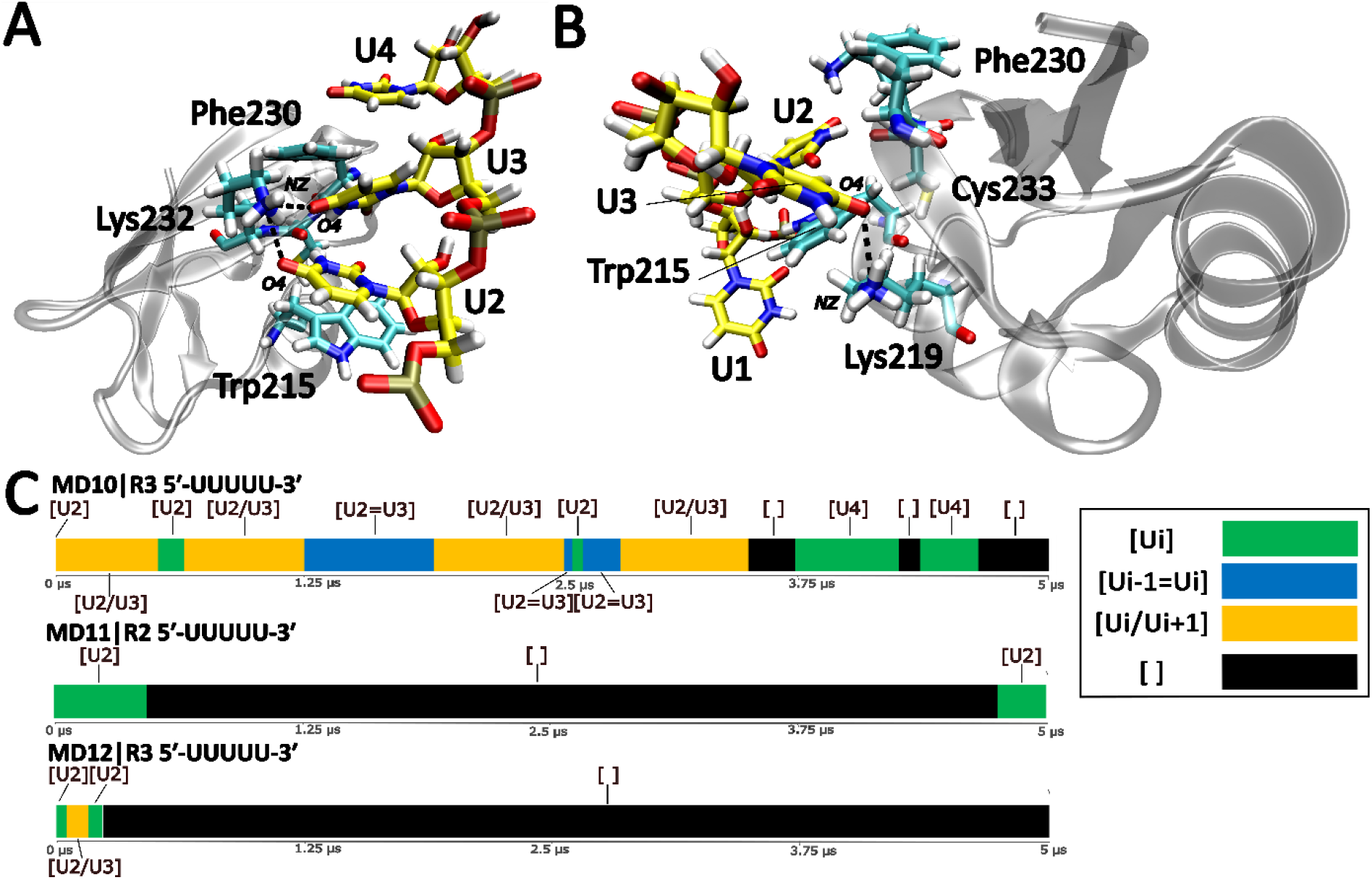
Role of the Lys232 and Lys219 residues vicinal to the binding pocket. (A) H-bonds formed between Lys232 and the uridines in the vertical state. (B) H-bond between the Ui+1 nucleotide in the horizontal state and Lys219. For clarity, not all the nucleotides are displayed. (C) Time-development of the binding pocket occupancy in selected simulations with mutated proteins. A single representative simulation of each system is shown (see the Supplementary Information for the rest).

On the other hand, simulations with the Lys219Ala mutant revealed behavior similar to the wild-type, except of slightly increased dynamics of the horizontal states and altered populations of some of the H-bonds (Supplementary Information). With both mutations combined, the same behavior as with the Lys232Ala substitution was observed.

### Spontaneous binding simulations detail the involvement of residues proximal to the binding pocket

We have carried out a set of simulations starting from an entirely unbound state (see Methods and ref. (23)). As observed earlier for other RRM-ssRNA complexes, (23) the RNA typically made a first contact with an arbitrary surface residue before later spontaneously moving towards the native binding site. As the RNA approached the binding area, the uridines formed stacking interactions with the residues proximal to the binding pocket (Figure 5, A and B) and Lys232 formed H-bonds and salt-bridges, guiding and pre-organizing the RNA binding. At the end of the binding process a single uridine would occupy the binding pocket (Figure 5, C). Subsequently, we observed the same equilibrium dynamics involving single uridine, horizontal, vertical and unbound states, as seen in the simulations started from the fully bound complexes (Figure 5, D). The protein also diffused along the RNA using the same mechanism as described above. We have observed more RNA dissociation events in our spontaneous binding simulations than in the simulations starting from the fully bound complex, a consequence of having used poly-(U) tetramer instead of pentamer (see Methods). This dissociation usually involved the RNA interacting with the residues proximal to the binding pocket. In other words, the uridines left the binding pocket, but the RNA maintained its molecular contact with the protein surface. More rarely, the RNA would completely depart from the protein surface. In both cases, new binding attempts typically followed. Lastly, note that in few of the binding simulations, the RNA did not manage to locate the binding site on the simulation timescale.

**Figure 5:**
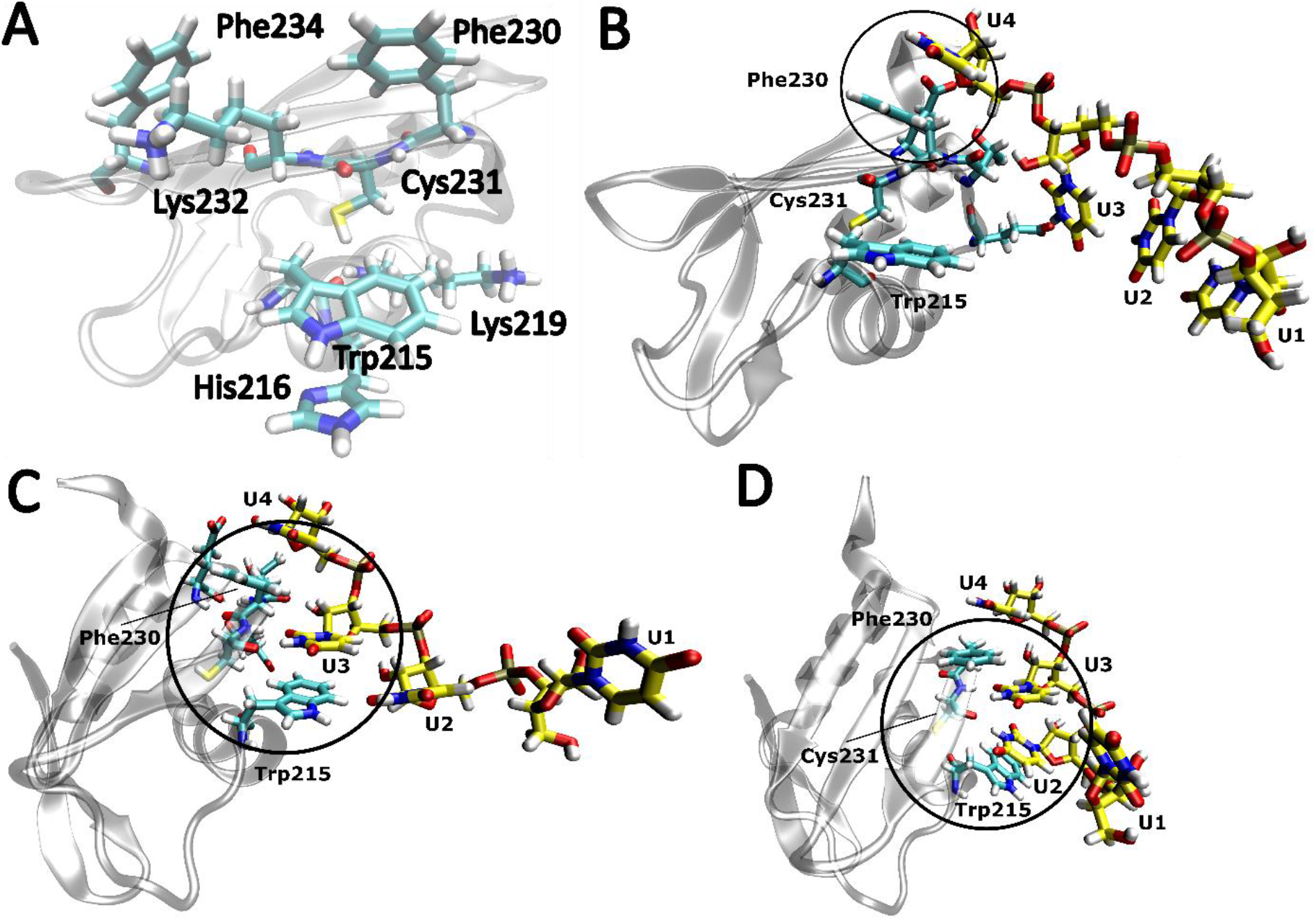
An example of the binding pathway of poly-(U) RNA to TbRGG2 RRM. Upon coming into proximity of the protein residues in vicinity of the binding pocket (A), stacking interactions formed between the RNA and these residues (B), shackling the RNA in this location and giving it time to gradually establish the native protein-RNA interface (C). (D) A vertical state formed later on.

## Discussion

### Equilibrium dynamics is integral to TbRGG2 RRM recognition of poly-(U) RNA

Here, we used unbiased MD simulations to explore the TbRGG2 RRM domain binding to its poly-(U) RNA substrate. We have carried out simulations starting from the experimentally determined (X-ray crystallography) structure as well as simulations starting from an entirely unbound state where the RNA and protein are separated. The results show good convergence, with qualitatively the same dynamics observed from both starting states. According to the experimental structure (37), the interface of the complex consists of a single binding pocket recognizing a single uridine. Such minimalistic interface is rather unusual among the RRM domains, which typically contain multiple binding pockets, recognizing several consecutive nucleotides (3). It was suggested that homodimerization of two TbRGG2 RRM domains compensates for its minimalistic interface (37). However, even two binding pockets selective for uridines and acting in tandem seem at first sight unlikely to achieve the necessary specificity and affinity which is in a micromolar range. Based on the MD simulations we suggest that TbRGG2 RRM utilizes extensive equilibrium dynamics to compensate for its solitary binding pocket. First, the simulations clearly confirm that TbRGG2 RRM contains just one binding site specifically recognizing uridines. The binding site is well reproduced in fully bound simulations, showing the uridine forming base-specific interactions with the protein backbone, with the base sandwiched between Trp215 and Phe230 aromatic side-chains (Figure 1). This binding site can be located by the RNA even in the spontaneous binding simulations. Several amino acids proximal to the binding site were found to be engaged in the binding process, helping to navigate the RNA to the native position and stabilizing the temporarily unbound states (Figure 5). Note that for fast initial binding it could be advantageous to have just one binding pocket as proteins with several pockets are more sensitive to the precise positioning of the incoming RNA and can have multiple binding registers that need to be explored (23). In general, the protein-RNA complexes represent a carefully orchestrated evolutionary compromise between affinity, specificity and the substrate exchange rates. The individual proteins need to be able to unambiguously recognize their specific target sequences in the vast pool of cellular RNAs (specificity), and they need to be able to hold onto them for a sufficient amount of time (affinity) to accomplish their biological function. An often underappreciated factor is that the proteins also need to be able to achieve all this on biologically relevant timescales, necessitating a swift discarding of non-cognate and semi-cognate RNAs encountered in the pool. The inevitable presence of numerous near-cognate RNA sequences limits how strong the protein-RNA affinity can get before slow substrate exchange starts hampering its biological purpose. Even once the protein encounters the target RNA, it needs to locate its specific cognate binding motif among potentially hundreds of nucleotides.

### Unique equilibrium dynamics of the TbRGG2 RRM complex effectively circumvents limitations of stochastic diffusion

The simulations of TbRGG2 RRM reveal a unique structural mechanism through which all the contradictory requirements placed on protein-RNA complexes can be elegantly reconciled and balanced. Namely, in addition to the single uridine binding state (shown by experiments) we have observed two additional binding states with two uridines simultaneously occupying the binding pocket, referring to these states as vertical and horizontal states (Figure 1). Formation of these additional states is facilitated by only a modest widening of the binding pocket by up to ∼1.0 Å. All three states are engaged in fast μs-scale interconversion dynamics. The RNA can also temporarily unbind from the binding pocket, transiently interacting with the residues proximal to the binding pocket before re-entering the pocket. More rarely, the RNA left the protein surface, which was followed by a new binding attempt. The interconversion dynamics among the three principal binding states allows a smooth exchange of the uridines in the binding pocket, facilitating a low-barrier and virtually unobstructed diffusion of the RRM along the whole poly-uridine sequence. Such recognition mechanism is uniquely suited for a solitary binding pocket as more pockets would require the bound RNA to shift its position in all pockets in a concerted manner. When lacking external energy provided, e.g., by ATP hydrolysis, such RNA shifts are accomplished strictly by stochastic movements (55). The traditional models of purely stochastic diffusion envision the protein step-by-step shifting along the nucleic acid in a linear fashion or repeatedly binding and unbinding at different sites set apart by few to hundreds of nucleotides, until reaching the cognate binding motif. However, strict application of both of these models leads to biologically unsustainable rates of exchange and recognition (56). Presence of a single binding pocket considerably lowers the energy barriers and elegantly circumvents the inherent limitations of stochastic diffusion.

In summary, our unbiased MD simulations indicate that the free-energy landscape of the bound TbRGG2–poly(U) complex is very broad (flat) and consists of a set of shallow states (or substates) separated by small free-energy barriers so that they interconvert on the μs time scale. Such structure of the free-energy surface allows fast movements of TbRGG2 RRM along the whole poly(U) ssRNA sequence and may contribute entropically to the binding affinity. We suggest the entire poly-(U) stretch where the RRM is bound essentially corresponds to a single continuous binding region at 300 K (Figure 6). Upon encountering non-native nucleotides like cytosines, TbRGG2 RRM is unable to stably accommodate them via any of the three binding states observed (Figure 1), causing the protein to revert back to the uridine-rich regions. Alternatively, it can “skip” the cytidine by the RNA temporarily leaving the binding pocket and rebinding further up-or downstream. In this fashion, TbRGG2 RRM is supremely tuned for rapidly locating the most U-rich region of any RNA sequence it encounters.

**Figure 6:**
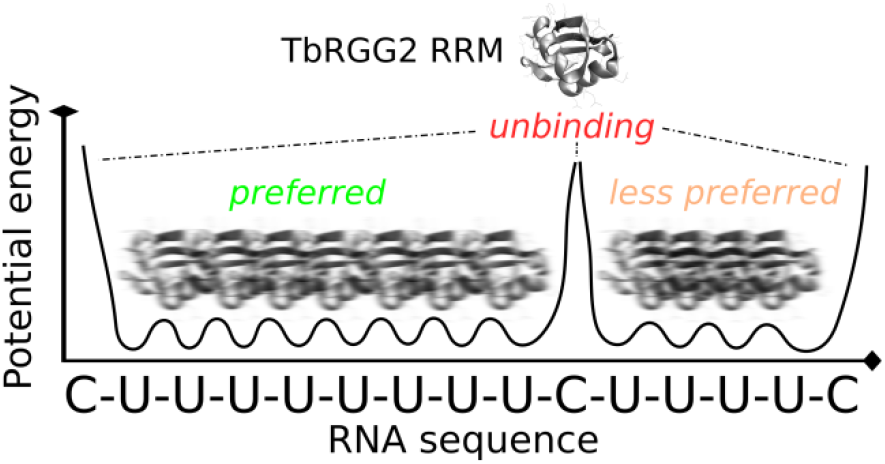
Scheme of the suggested equilibrium dynamics of TbRGG2 RRM bound to U-rich RNA. The flexible binding pocket allows low-barrier transitions along the continuous poly-(U) regions of the RNA on the microsecond timescale at physiological temperatures, where it will essentially form a single flat mildly undulated basin on the potential energy surface. The fuzzy protein images above the individual poly-(U) stretches indicate the protein rapidly shifting within the region. Meanwhile, encounter with non-uridine nucleotide destabilizes the complex and can lead to unbinding and another binding attempt. We suggest this mechanism allows the protein to quickly locate the longest stretches of uridines which then become the preferred binding sites by entropic effects. Utilization of only a single binding pocket ensures non-cognate substrates are rapidly discarded while the entropically advantageous equilibrium dynamics accessible only for the cognate sequence guarantees high specificity. The one-dimensional potential energy plot is merely illustrative, being deduced from the simulation analyses; it is not a result of any direct measurements or computations.

### Spontaneous binding simulations demonstrate the role of the residues proximal to the binding pocket and confirm the unique equilibrium dynamics

After a random first contact with any part of the protein surface, the binding process commenced with the RNA approaching the hydrophobic or positively charged residues proximal to the binding pocket (Figure 5A), forming non-specific interactions with them. In simulations started from a fully bound complex, the same residues would bind the RNA when it temporarily left the binding pocket. The protein segments surrounding the binding site might be guiding or funnelling the complex towards its free energy minimum. Such non-specific contacts are similar in principle to the “pre-binding state” observed for the HuR RRM3 (23), although less structurally defined. Upon formation of native binding, our spontaneous binding simulations showed qualitatively identical equilibrium dynamics as the simulations started from fully bound complex, confirming the results are not overly dependent on the starting structure. At the same time, the spontaneous binding simulations suggested that the vertical and horizontal binding modes are not part of the binding-unbinding landscape of the complex as no binding events via these substates were observed. Instead, they appear to be distinct conformational states associated with the low-barrier equilibrium dynamics of the fully bound complex.

Lastly, we have also attempted spontaneous binding simulations of the poly-(G) RNA. The poly-(G) sequence is known to be bound by TbRGG2 RRM through a binding site distinct from the poly-(U) (37). However, the bound structure showing the poly-(G) RNA binding is not available. Unfortunately, we were unable to make any conclusions from our spontaneous binding simulations either (not shown). We speculate the binding of poly-(G) sequence may involve some hitherto unknown conformational change of the protein that was not sampled on our simulation timescale.

### Dimerization of TbRGG2 RRM combined with its equilibrium dynamics could further enhance the selectivity for U-rich RNA sequences

TbRGG2 possesses a dimerization domain located just C-terminal of the RRM (37). However, the dimerization domain has not been included in the experimentally determined X-ray structure of the RRM (PDB: 6E4P), and while the RRMs are making molecular contacts in the asymmetric unit, it is not directly stated that these correspond to the native dimers (37). In fact it is unlikely, as the relative positions of the RRMs within the crystal lattice do not allow straightforward addition of the dimerization domains by molecular modeling. Furthermore, our simulations of the putative RRM homodimers found in the crystal lattice, both with and without the RNA bound, led to almost instantaneous and permanent loss of the experimental protein-protein interface (Supplementary Information). The RRMs remained in some molecular contact but failed to establish any interface that would be stable (Supplementary Information). Even in absence of the C-terminal dimerization domains, such swift loss of native interface interactions should not occur in MD simulations for even very weakly bound dimers, especially considering that the *ff* generally overstabilizes the solute-solute interactions (23). We therefore conclude the arrangement and interactions of the TbRGG2 RRMs as observed in the experimental structure reflects the constraints of the crystal lattice rather than a preference for a particular dimer interface. Multiple protein chains tightly surrounding and binding the single RNA molecule might also be what had stalled the RNA diffusion mechanism and allowed structure resolution. This suggests the TbRGG2 homodimerization is driven solely by the C-terminal domains, placing the RRMs in close proximity but with no major direct interactions between them. The equilibrium dynamics of the RRM-ssRNA interface observed in this work can then be expected to independently occur for both binding pockets of the two RRMs also within the homodimer context of full length TbRGG2. With two RRMs acting like independent “walkers” along the RNA sequence, the specificity of TbRGG2 for U-rich parts of the RNA sequences would be vastly increased.

## Conclusions

Using large-scale unbiased atomistic molecular dynamics simulations, we showed that the TbRGG2 RRM binding site for poly-(U) RNAs is characterized by low-barrier interconversion dynamics involving three distinct binding substates, allowing a smooth exchange of the uridines in the binding pocket. All three binding substates are highly selective for uridines and we observed completely spontaneous diffusion of the poly-(U) sequences all the way to the strand termini. Presence of non-native cytidines tends to cause either swift transition back towards the U-rich regions or eventual disengagement of the complex followed by another binding attempt elsewhere on the sequence. Using this mechanism, TbRGG2 RRM should be able to rapidly locate the most uridine-rich part of any RNA sequence. Spontaneous binding simulations initiated from the two molecules fully separated reveal qualitatively identical behaviour, confirming the observed equilibrium dynamics of the fully bound complex is not part of the binding-unbinding dynamics. Dynamical binding mechanisms and sites such as the one employed by TbRGG2 RRM could constitute a so far under-appreciated element of protein-RNA recognition, difficult to detect by conventional structural biology experiments, but with potentially high utility in the astonishingly diverse world of protein-RNA complexes.

## Supplementary Information

Additional information on Stafix; simulations of the putative TbRGG2 RRM homodimers; additional information on the intermolecular H-bonds formed by single; vertical and horizontal binding states; comments on the binding states formed by cytidines. Supplementary Figures, Graphs and Tables.

## Acknowledgements

This work has been conducted in the sustainability period of the project SYMBIT No. CZ.02.1.01/0.0/0.0/15_003/0000477 as its follow-up activity.

