## Supplementary for "How Binding Site Flexibility Promotes RNA Scanning in TbRGG2 RRM: A Molecular Dynamics Simulation Study"

### Supplementary Information Text

**Additional comments on the use of Stafix in TbRGG2 RRM simulations.**

The standard OL3 RNA force field (*ff*) overestimates the intra-RNA interactions, causing overcompaction of the simulated ssRNAs (1). In context of protein-RNA complexes, the overstabilized RNA-RNA interactions eventually outcompete even the native intermolecular interfaces, leading to complex destabilization. Swiftness of this destabilization varies from system to system, but it is generally quick in complexes with very dynamic interfaces. We recently developed a system-specific *ff* correction called ***Stafix***, which suppresses this undesired behavior, allowing us to observe completely spontaneous binding of unstructured ssRNAs to proteins while also greatly improving simulations of fully bound protein-RNA complexes (1). In case of the 5′-UUUUU-3′ RNA (the most commonly simulated RNA in this study), Stafix increases the average solvent-accessible surface of such RNA by ~12% (1), allowing spontaneous capture by proteins and preventing loss of the fully formed complexes due to RNA overcompaction.

**Instability of the TbRGG2 RRM dimer.**

Biochemical experiments indicated the TbRGG2 protein forms a homodimer in solution via a dimerization domain located just C-terminal of the RRM. The dimerization domain was not present in the crystallization experiments along with the RRM, however, the X-ray structures deposited in the database contain multiple RRMs in close molecular contact within the asymmetric unit. This raises the possibility some of these RRMs could correspond to the native homodimer, even when lacking the dimerization domain. However, they could also be ordinary crystal packing interactions. We note the authors of the paper describing the structures do not directly state whether the presented structures correspond to the native TbRGG2 RRM dimer (2). Indeed, visual inspection of the structures did not reveal many intermolecular interactions typically seen with stable protein dimers. To make more definitive conclusions, we have decided to carry out simulations of the two potential TbRGG2 RRM homodimer arrangements as observed in the crystal lattice (MDS1 and MDS2; see Table S1) to assess their stability. In both systems, we observed almost immediate loss of the original molecular interface between the two RRMs (Figure S4). The RRMs remained in contact and the system subsequently explored various interactions and mutual orientations of the two

RRMs, but failed to settle on any specific dimer interface. We then performed additional simulations of the more stable (but still very unstable) of the two putative homodimers with the RNA bound (MDS3), to see whether the RNA presence could potentially stabilize the structure. However, we observed the same swift loss of the molecular interface between the two RRMs, suggesting the bound RNA does not serve as a stabilizing element for the putative homodimer. Based on these results, we conclude the dimerization of TbRGG2, as observed in biochemical experiments (2), is almost certainly driven solely by the C-terminal dimerization domain, with limited direct molecular interaction between the RRMs. Were such direct RRM-RRM contacts structurally significant, we would expect such dimers to be stable in MD simulations even without the dimerization domain, at least for a short time. Alternatively, the simulation should have been able to locate the native interface after the one in the starting structure became lost. In conclusion, the virtually immediate disintegration of the experimentally observed protein-protein interface suggests that the experimental structure of the TbRGG2 RRM does not display the native dimer structure but rather common crystal packing interactions (2). Lastly, we note the isolated RRM domain (with the RNA removed) was fully stable in simulations (MDS4; see Table S1), with no significant changes compared to the starting structure observed.

**Intermolecular H-bonds observed in the single, vertical and horizontal states of the binding pocket.**

The populations of intermolecular H-bonds formed by TbRGG2 RRM complex are influenced by the current state of the binding pocket (Figure S2). Here we provide a more detailed description of the intermolecular H-bond patterns characteristic for the single uridine, vertical, and horizontal states (see the main text Figure 1). The analyses were carried out by observing all three states in the ensemble of the MD2, MD3 and MD4 simulations and the populations of the individual H-bonds are listed in Tables S2, S3 and S4. In addition to the principal Ui(N3)-Cys231(O) H-bond (see the main text), the ***single uridine state*** was also characterized by the Ui(O2)-Cys231(N) H-bond (Table S2 and Figure S2A). There was also the Thr229(O)-Ui(O2′) interaction which aided the binding by lowering the mobility of the sugar ring and aligning the Ui base for proper interaction with the Cys231. Lastly, the Lys232 side-chain formed transient H-bonds with RNA in most of the replicates. In the ***vertical state***, in addition to the bifurcated H-bond between the uridine N3 atoms and the Cys231(O) (see the main text), the uridine closer to Phe230 formed Ui(O2)-Cys231(N) H-bond while the uridine closer to Trp215 formed H-bonds with Val233 (Table S3). In the ***horizontal state***, the Ui+1 nucleotide (main text Figure 1) made contact with the hydrophobic patch (Figure S1) and also formed intermolecular H-bonds with Val228 and Lys219 (Table S4). The nucleotide deeper in the binding pocket (Ui) formed similar H-bonds as the nucleotides of the vertical state closer to Trp215. Apart from the intermolecular H-bonds, intramolecular RNA H-bonds were also observed in all three states, contributing to the proper orientation of the RNA molecule towards the RRM. The most prevalent was the Ui+1:O5′ - Ui:O2′ H-bond, with occupancies sometimes over 50%.

**The diverse binding states observed in simulations with cytidine-containing RNA**

As noted in the main text, TbRGG2 RRM tends to explore numerous arrangements of the binding pocket when cytidine is present, in an attempt to stabilize such inherently unstable interface. This is especially true for the “Ci-∞” conformations in which both uridine and cytosine are present in the binding pocket (Figure 3B and 3D). Most common of those is the *hybrid state* (see the main text). Other arrangements mainly observed in the MD5 simulations included the uridine oriented parallel or askew to Trp215 and the cytidine slightly shifted towards the hydrophobic patch which normally stabilizes the horizontal state in poly-(U)

RNA binding (Figure 1C; Figure 2). Another arrangement, mostly observed in the MD8 simulations, had the uridine sandwiched between cytosine and Trp215 while the backbone between nucleotides C3 and C4 sporadically interacted with the binding pocket. Lastly, we note that there were sometimes prolonged instances of single cytidine binding (labeled as [Ci]). However, these were commonly supported by interactions with the RNA backbone (Figure 3, D) in addition to the fluctuating H-bonds Cys231(O)-Ci(N4) and Ci(O2)-Val233(N). In conclusion, all instances of cytidine binding were greatly unstable compared to the uridine binding. The greater variety of conformational states explored by the system reflects the attempts (and failures) of TbRGG2 RRM to somehow establish stable binding.

### Supplementary Information Figures

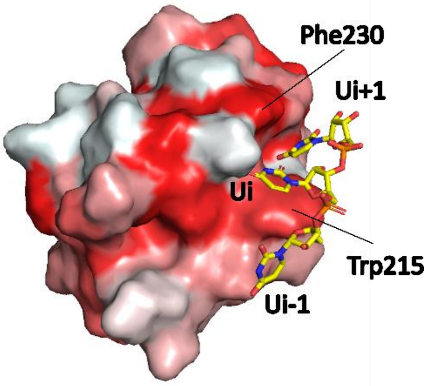


Figure S1: **Visualization of the hydrophobic patch stabilizing the horizontal state.** A) The protein surface is colored according to the Eisenberg hydrophobicity scale (3), with red regions more hydrophobic compared to white regions. The RNA is shown in the horizontal state [Ui/Ui+1] where Ui+1 is encircled by the hydrophobic patch, formed by surface of Trp215, the aliphatic part of Lys219 and the side chain of Val228.

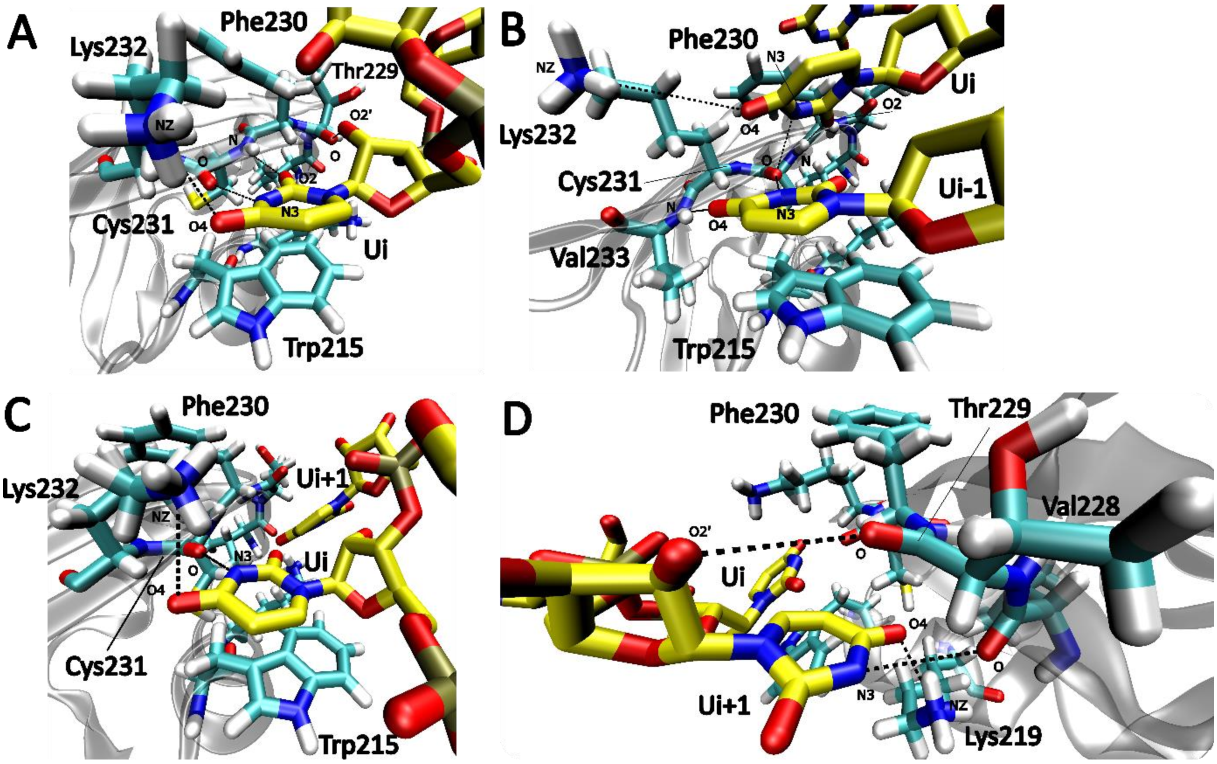


Figure S2: **Visualization of the major intermolecular H-bonds formed by the binding pocket.** (A) The single uridine state with H-bonds Ui(O4)–Lys232(NZ), Cys231(O)–Ui(N3), Ui(O2)–Cys231(N) and Thr229(O)–Ui(O2′); (B) The vertical state with H-bonds Ui(O4)–Lys232(NZ), Ui-1(O4)–Val233(N), Cys231(O)–Ui-1(N3), Cys231(O)–Ui(N3) and Ui(O2)–Cys231(N); (C) Ui view of the horizontal state with H-bonds U2(O4) - Lys232(NZ) and Cys231(O) – U2(N3); (D) Ui+1 view of the horizontal state with H-bonds Thr229(O) – Ui+1(O2′), Val228(O) – Ui+1(N3) and Ui+1(O4) – Lys219(NZ). All H-bonds are listed from left to right. Note that the uridines are labeled in analogy with main text Figure 1.

| | | |
|---|---|---|
| **[Ui]** | | ***Single uridine state*** - the binding pocket occupied by one uridine (main text Figure 1A). |
| **[Ci]** | | The binding pocket occupied by one cytidine (main text Figure 3A). |
| **[Ui-1=Ui]** | | ***Vertical state*** - the binding pocket occupied by two consecutive uridines with their bases stacked. Ui-1 and Ui are closer to Trp215 and Phe230, respectively (see the main text Figure 1B). |
| **[Ui/Ui+1]** | | ***Horizontal state*** – the binding pocket occupied by two consecutive uridines Ui and Ui+1. Their bases are positioned in the same plane and the latter is stabilized by the hydrophobic patch (Figure 1C and Figure S1). |
| **[Ci-∞]** | | Binding pocket occupied by at least one cytidine and another base, in any conformational arrangement. |
| **[Ui=Uj] or [Ui/Uj]** | | Vertical or horizontal states formed by non-consecutive residues (\|i-j\|>1). |
| **[ ]** | | No nucleotide within the binding pocket. |

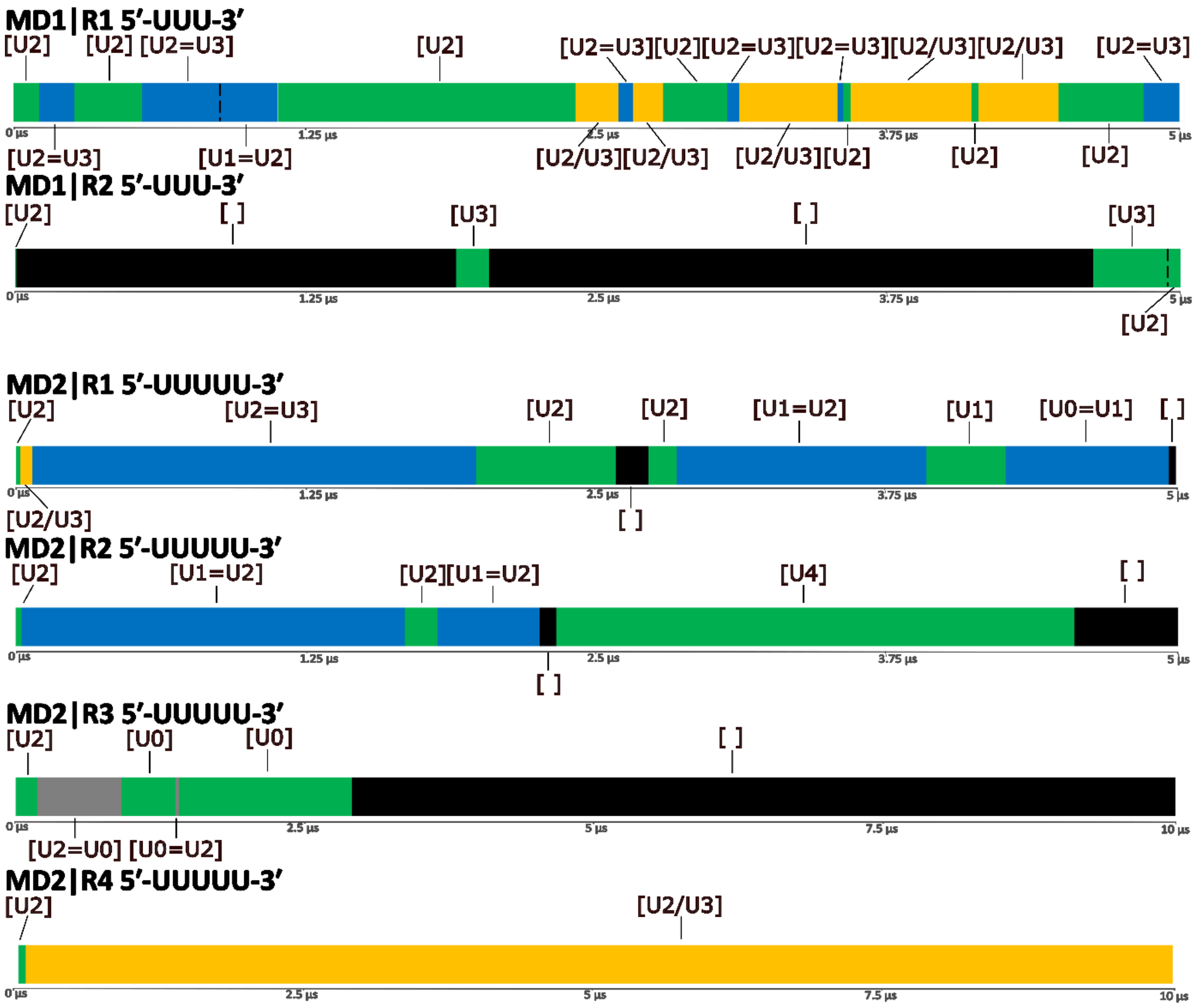

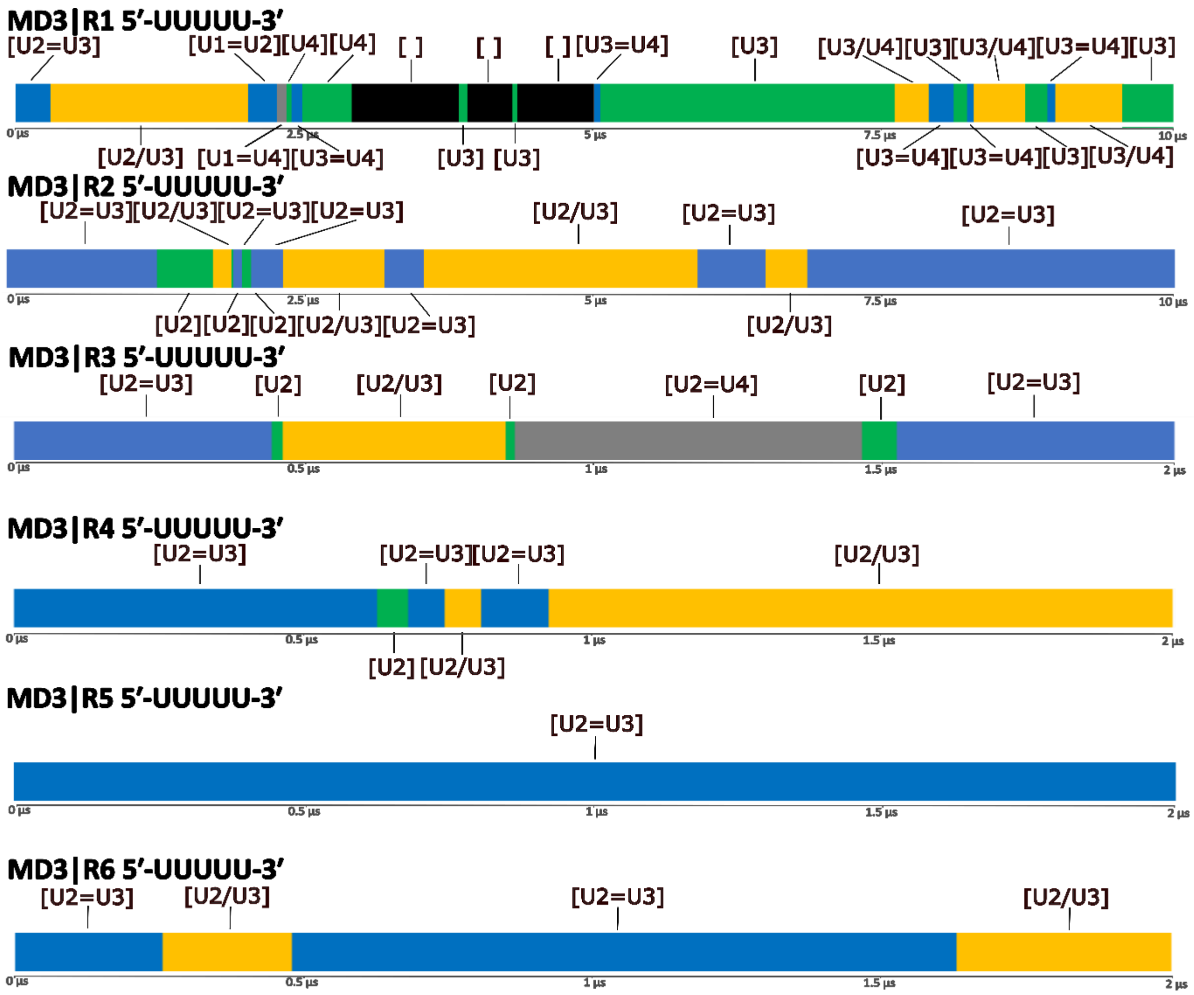
MD3|R1 5′-UUUUU-3′
[U2=U3]
[U1=U2][U4][U4]
[ ]
[ ]
[ ] [U3=U4]
[U3]
[U3/U4][U3][U3/U4][U3=U4][U3]
0 µs
2.5 µs
5 µs
7.5 µs
10 µs
[U2/U3] [U1=U4][U3=U4]
[U3] [U3]
[U3=U4][U3=U4][U3][U3/U4]
MD3|R2 5′-UUUUU-3′
[U2=U3][U2/U3][U2=U3][U2=U3]
[U2/U3]
[U2=U3]
[U2=U3]
0 µs
2.5 µs
5 µs
7.5 µs
10 µs
[U2][U2][U2][U2/U3][U2=U3]
[U2/U3]
MD3|R3 5′-UUUUU-3′
[U2=U3]
[U2]
[U2/U3]
[U2]
[U2=U4]
[U2]
[U2=U3]
0 µs
0.5 µs
1 µs
1.5 µs
2 µs
MD3|R4 5′-UUUUU-3′
[U2=U3]
[U2=U3][U2=U3]
[U2/U3]
0 µs
0.5 µs
1 µs
1.5 µs
2 µs
[U2] [U2/U3]
MD3|R5 5′-UUUUU-3′
[U2=U3]
0 µs
0.5 µs
1 µs
1.5 µs
2 µs
MD3|R6 5′-UUUUU-3′
[U2=U3]
[U2/U3]
[U2=U3]
[U2/U3]
0 µs
0.5 µs
1 µs
1.5 µs
2 µs

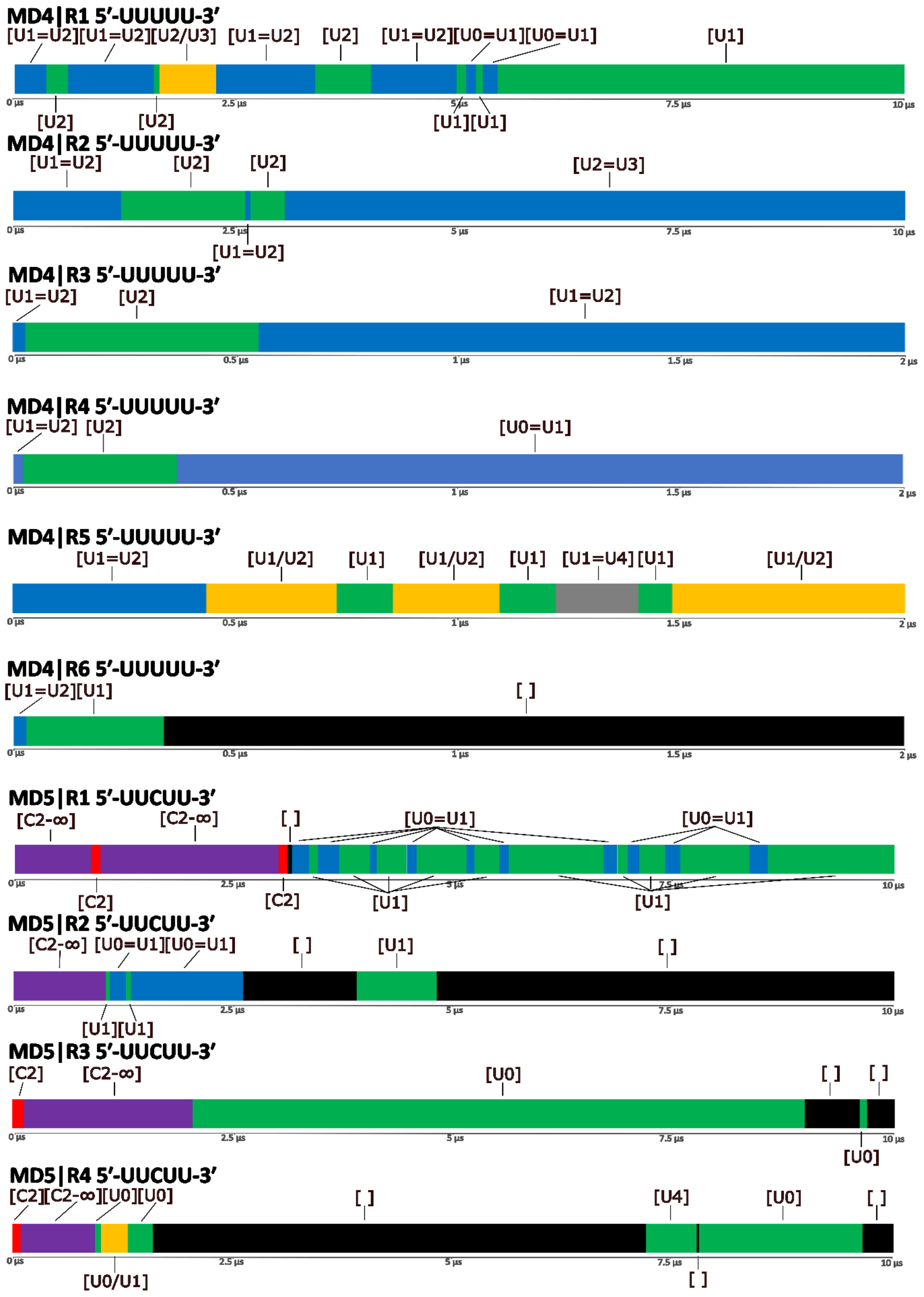

MD4|R1 5′-UUUUU-3′
[U1=U2][U1=U2][U2/U3] [U1=U2] [U2] [U1=U2][U0=U1][U0=U1] [U1]
0 µs 2.5 µs 5 µs 7.5 µs 10 µs
[U2] [U2] [U1][U1]
MD4|R2 5′-UUUUU-3′
[U1=U2] [U2] [U2] [U2=U3]
0 µs 2.5 µs 5 µs 7.5 µs 10 µs
[U1=U2]
MD4|R3 5′-UUUUU-3′
[U1=U2] [U2] [U1=U2]
0 µs 0.5 µs 1 µs 1.5 µs 2 µs
MD4|R4 5′-UUUUU-3′
[U1=U2] [U2] [U0=U1]
0 µs 0.5 µs 1 µs 1.5 µs 2 µs
MD4|R5 5′-UUUUU-3′
[U1=U2] [U1/U2] [U1] [U1/U2] [U1] [U1=U4] [U1] [U1/U2]
0 µs 0.5 µs 1 µs 1.5 µs 2 µs
MD4|R6 5′-UUUUU-3′
[U1=U2][U1] [ ]
0 µs 0.5 µs 1 µs 1.5 µs 2 µs
MD5|R1 5′-UUCUU-3′
[C2-∞] [C2-∞] [ ] [U0=U1] [U0=U1]
0 µs 2.5 µs 5 µs 7.5 µs 10 µs
[C2] [C2] [U1] [U1]
MD5|R2 5′-UUCUU-3′
[C2-∞] [U0=U1][U0=U1] [ ] [U1] [ ]
0 µs 2.5 µs 5 µs 7.5 µs 10 µs
[U1][U1]
MD5|R3 5′-UUCUU-3′
[C2] [C2-∞] [U0] [ ] [ ]
0 µs 2.5 µs 5 µs 7.5 µs 10 µs
[U0]
MD5|R4 5′-UUCUU-3′
[C2][C2-∞][U0][U0] [ ] [U4] [U0] [ ]
0 µs 2.5 µs 5 µs 7.5 µs 10 µs
[U0/U1] [ ]

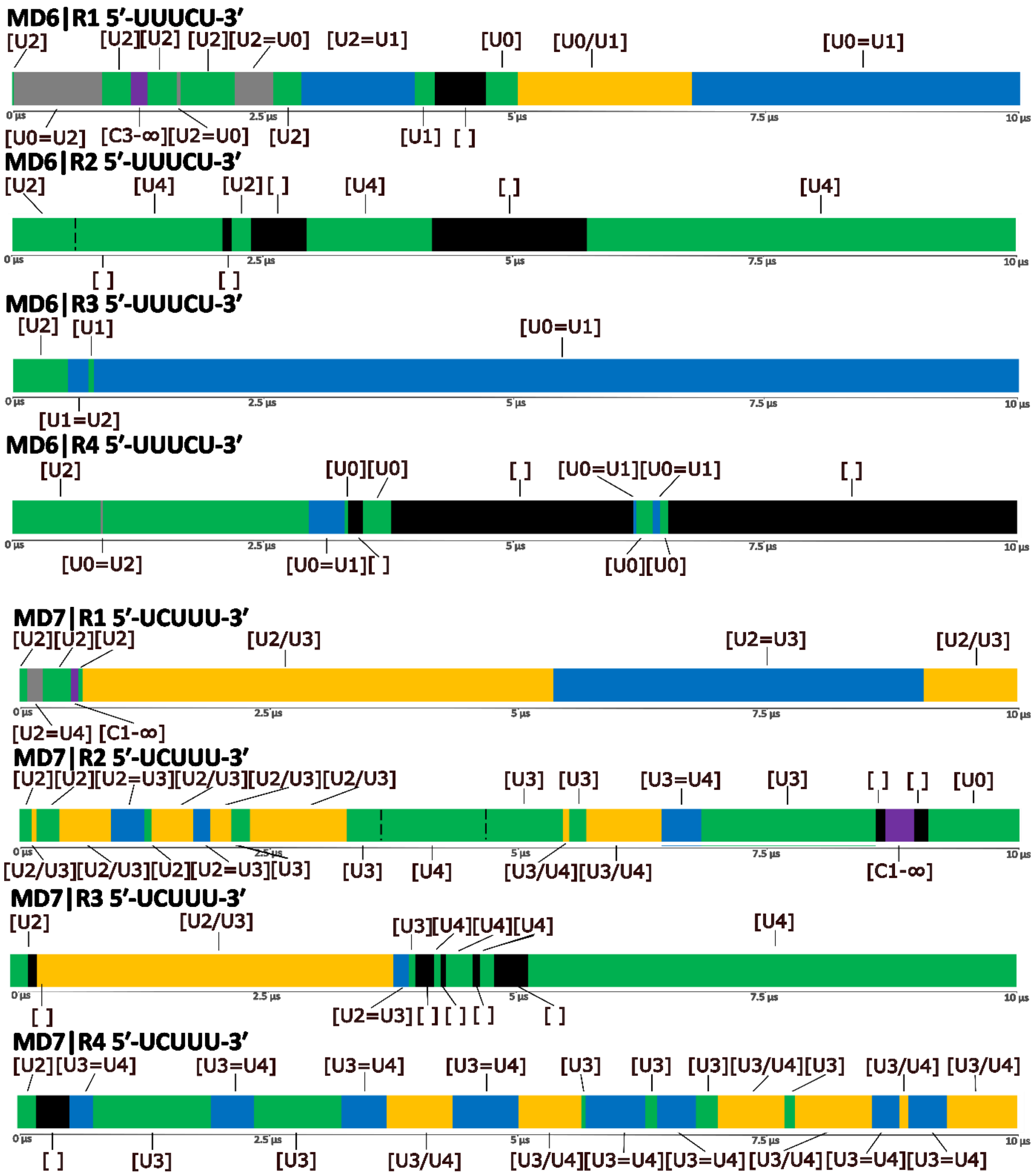
MD6|R1 5′-UUUCU-3′
[U2]
[U2][U2] [U2][U2=U0] [U2=U1]
[U0]
[U0/U1]
[U0=U1]
0 µs
2.5 µs
5 µs
7.5 µs
10 µs
[U0=U2] [C3-∞][U2=U0] [U2]
[U1] [ ]
MD6|R2 5′-UUUCU-3′
[U2]
[U4]
[U2][ ]
[U4]
[ ]
[U4]
0 µs
2.5 µs
5 µs
7.5 µs
10 µs
[ ]
[ ]
MD6|R3 5′-UUUCU-3′
[U2] [U1]
[U0=U1]
0 µs
2.5 µs
5 µs
7.5 µs
10 µs
[U1=U2]
MD6|R4 5′-UUUCU-3′
[U2]
[U0][U0]
[ ]
[U0=U1][U0=U1]
[ ]
0 µs
2.5 µs
5 µs
7.5 µs
10 µs
[U0=U2]
[U0=U1][ ]
[U0][U0]
MD7|R1 5′-UCUUU-3′
[U2][U2][U2]
[U2/U3]
[U2=U3]
[U2/U3]
0 µs
2.5 µs
5 µs
7.5 µs
10 µs
[U2=U4] [C1-∞]
MD7|R2 5′-UCUUU-3′
[U2][U2][U2=U3][U2/U3][U2/U3][U2/U3]
[U3] [U3]
[U3=U4]
[U3]
[ ] [ ]
[U0]
0 µs
2.5 µs
5 µs
7.5 µs
10 µs
[U2/U3][U2/U3][U2][U2=U3][U3]
[U3]
[U4]
[U3/U4][U3/U4]
[C1-∞]
MD7|R3 5′-UCUUU-3′
[U2]
[U2/U3]
[U3][U4][U4][U4]
[U4]
0 µs
2.5 µs
5 µs
7.5 µs
10 µs
[ ]
[U2=U3][ ] [ ] [ ]
[ ]
MD7|R4 5′-UCUUU-3′
[U2][U3=U4]
[U3=U4]
[U3=U4]
[U3=U4]
[U3]
[U3]
[U3][U3/U4][U3]
[U3/U4] [U3/U4]
0 µs
2.5 µs
5 µs
7.5 µs
10 µs
[ ]
[U3]
[U3]
[U3/U4]
[U3/U4][U3=U4][U3=U4][U3/U4][U3=U4][U3=U4]

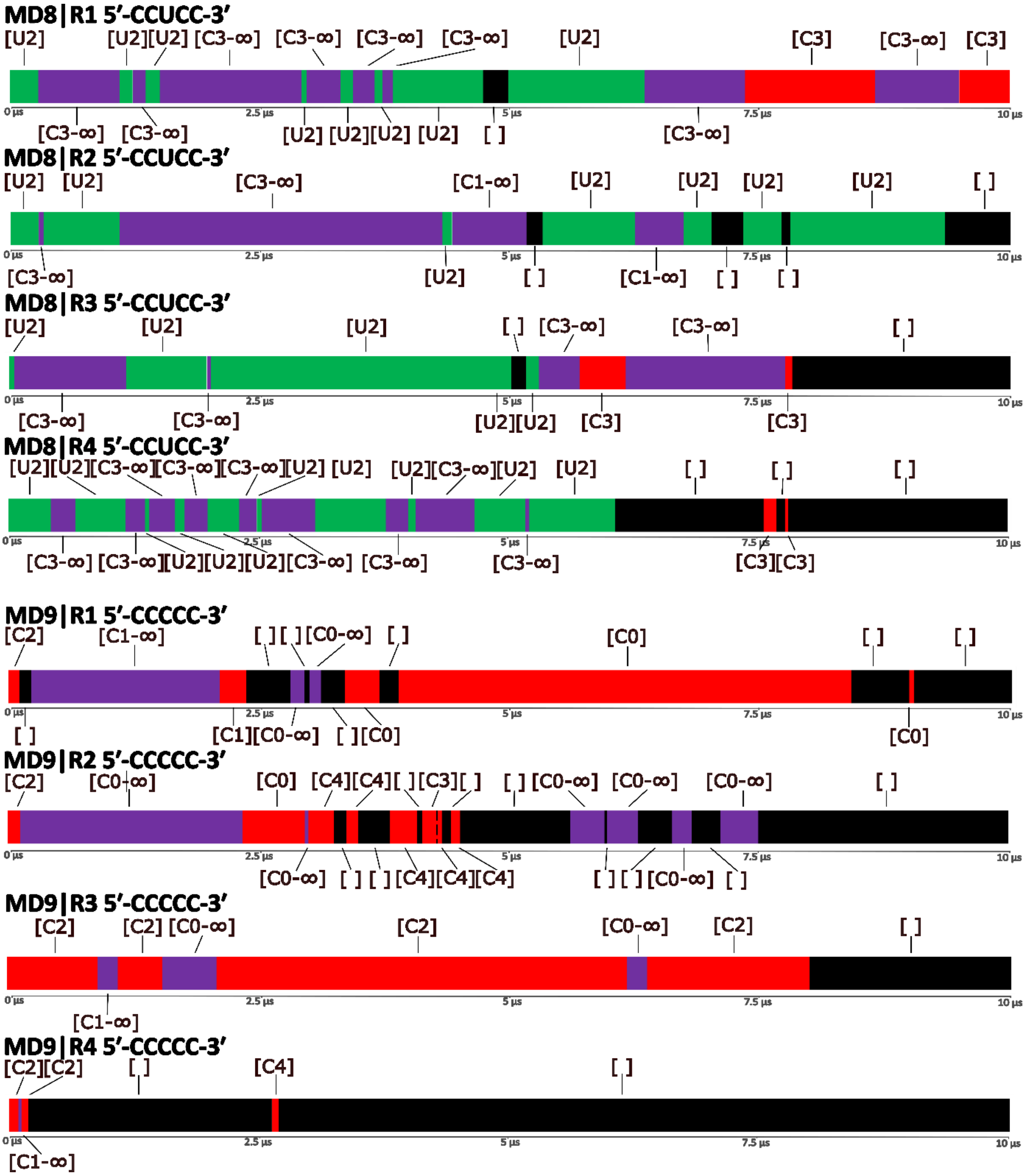
MD8|R1 5′-CCUCC-3′
[U2] [U2][U2] [C3-∞] [C3-∞] [C3-∞] [C3-∞] [U2] [C3] [C3-∞] [C3]
[C3-∞] [C3-∞] [U2] [U2][U2] [U2] [ ] [C3-∞]
MD8|R2 5′-CCUCC-3′
[U2] [U2] [C3-∞] [C1-∞] [U2] [U2] [U2] [U2] [ ]
[C3-∞] [U2] [ ] [C1-∞] [ ] [ ]
MD8|R3 5′-CCUCC-3′
[U2] [U2] [U2] [ ] [C3-∞] [C3-∞] [ ]
[C3-∞] [C3-∞] [U2][U2] [C3] [C3]
MD8|R4 5′-CCUCC-3′
[U2][U2][C3-∞][C3-∞][C3-∞][U2] [U2] [U2][C3-∞][U2] [U2] [ ] [ ] [ ]
[C3-∞] [C3-∞][U2][U2][U2][C3-∞] [C3-∞] [C3-∞] [C3][C3]
MD9|R1 5′-CCCCC-3′
[C2] [C1-∞] [ ] [ ] [C0-∞] [ ] [C0] [ ] [ ]
[ ] [C1][C0-∞] [ ] [C0] [C0]
MD9|R2 5′-CCCCC-3′
[C2] [C0-∞] [C0] [C4][C4][ ] [C3][ ] [ ] [C0-∞] [C0-∞] [C0-∞] [ ]
[C0-∞] [ ] [ ] [C4][C4][C4] [ ] [ ] [C0-∞] [ ]
MD9|R3 5′-CCCCC-3′
[C2] [C2] [C0-∞] [C2] [C0-∞] [C2] [ ]
[C1-∞]
MD9|R4 5′-CCCCC-3′
[C2][C2] [ ] [C4] [ ]
[C1-∞]
0 µs
2.5 µs
5 µs
7.5 µs
10 µs

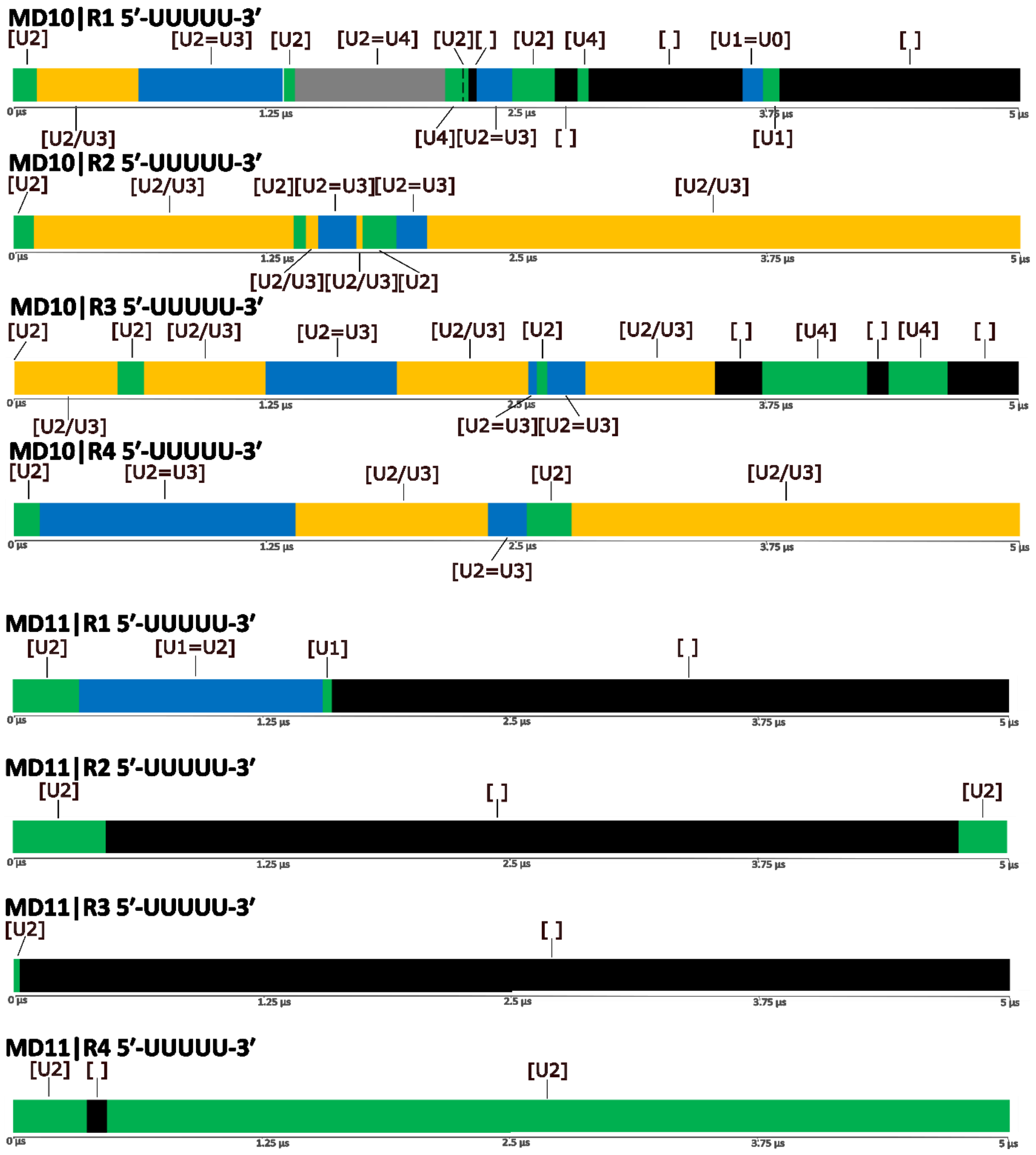
MD10|R1 5′-UUUUU-3′
[U2] [U2=U3] [U2] [U2=U4] [U2][ ] [U2] [U4] [ ] [U1=U0] [ ]
0 µs 1.25 µs 2.5 µs 3.75 µs 5 µs
[U2/U3] [U4][U2=U3] [ ] [U1]
MD10|R2 5′-UUUUU-3′
[U2] [U2/U3] [U2][U2=U3][U2=U3] [U2/U3]
0 µs 1.25 µs 2.5 µs 3.75 µs 5 µs
[U2/U3][U2/U3][U2]
MD10|R3 5′-UUUUU-3′
[U2] [U2] [U2/U3] [U2=U3] [U2/U3] [U2] [U2/U3] [ ] [U4] [ ] [U4] [ ]
0 µs 1.25 µs 2.5 µs 3.75 µs 5 µs
[U2/U3] [U2=U3][U2=U3]
MD10|R4 5′-UUUUU-3′
[U2] [U2=U3] [U2/U3] [U2] [U2/U3]
0 µs 1.25 µs 2.5 µs 3.75 µs 5 µs
[U2=U3]
MD11|R1 5′-UUUUU-3′
[U2] [U1=U2] [U1] [ ]
0 µs 1.25 µs 2.5 µs 3.75 µs 5 µs
MD11|R2 5′-UUUUU-3′
[U2] [ ] [U2]
0 µs 1.25 µs 2.5 µs 3.75 µs 5 µs
MD11|R3 5′-UUUUU-3′
[U2] [ ]
0 µs 1.25 µs 2.5 µs 3.75 µs 5 µs
MD11|R4 5′-UUUUU-3′
[U2] [ ] [U2]
0 µs 1.25 µs 2.5 µs 3.75 µs 5 µs

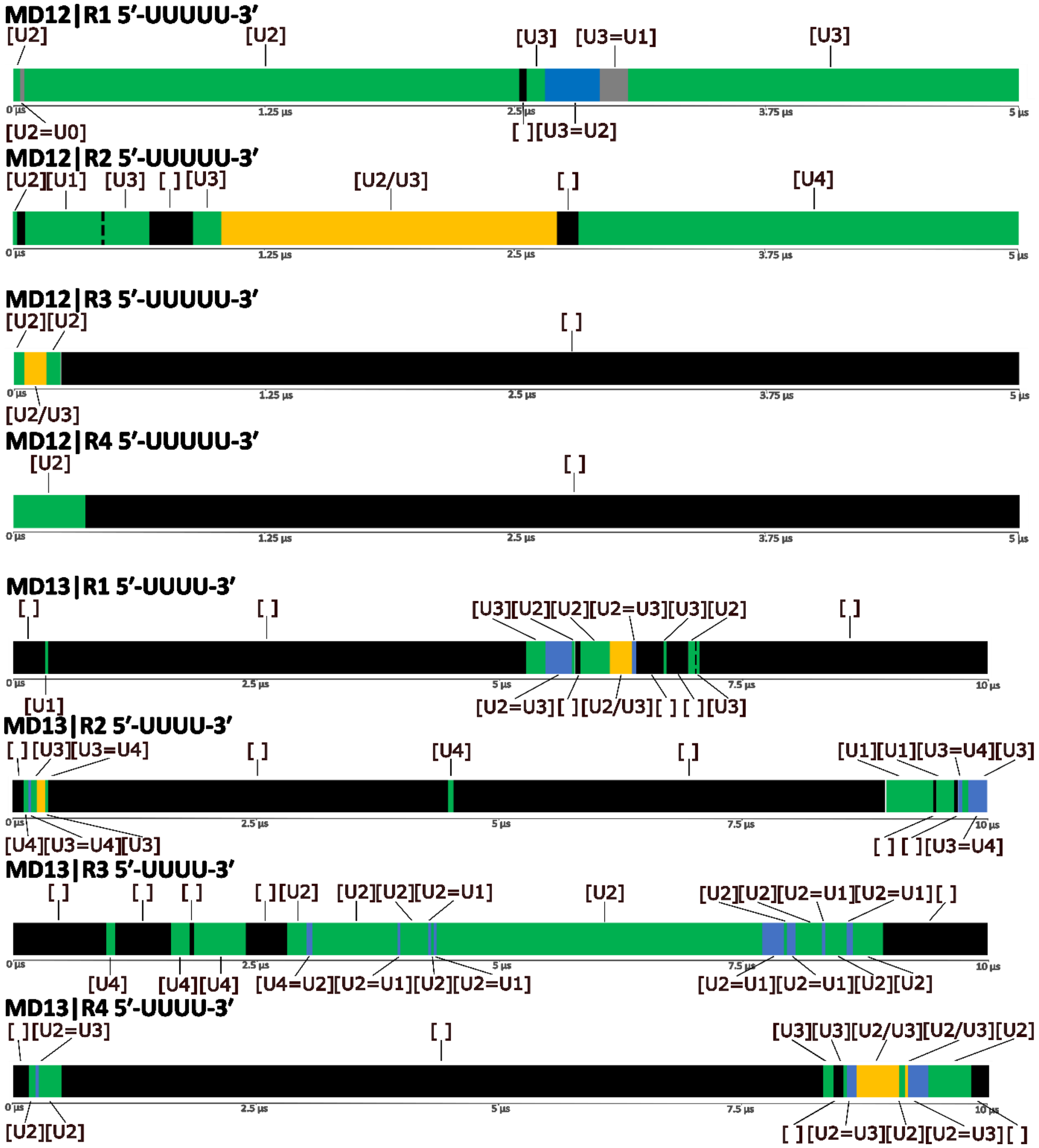
MD12|R1 5′-UUUUU-3′
[U2]
[U2]
[U3]
[U3=U1]
[U3]
0 µs
1.25 µs
2.5 µs
3.75 µs
5 µs
[U2=U0]
[ ][U3=U2]
MD12|R2 5′-UUUUU-3′
[U2][U1]
[U3]
[ ] [U3]
[U2/U3]
[ ]
[U4]
0 µs
1.25 µs
2.5 µs
3.75 µs
5 µs
MD12|R3 5′-UUUUU-3′
[U2][U2]
[ ]
0 µs
1.25 µs
2.5 µs
3.75 µs
5 µs
[U2/U3]
MD12|R4 5′-UUUUU-3′
[U2]
[ ]
0 µs
1.25 µs
2.5 µs
3.75 µs
5 µs
MD13|R1 5′-UUUU-3′
[ ]
[ ]
[U3][U2][U2][U2=U3][U3][U2]
[ ]
0 µs
2.5 µs
5 µs
7.5 µs
10 µs
[U1]
[U2=U3][ ][U2/U3][ ] [ ][U3]
MD13|R2 5′-UUUU-3′
[ ] [U3][U3=U4]
[ ]
[U4]
[ ]
[U1][U1][U3=U4][U3]
0 µs
2.5 µs
5 µs
7.5 µs
10 µs
[U4][U3=U4][U3]
[ ] [ ][U3=U4]
MD13|R3 5′-UUUU-3′
[ ]
[ ]
[ ]
[ ][U2]
[U2][U2][U2=U1]
[U2]
[U2][U2][U2=U1][U2=U1][ ]
0 µs
2.5 µs
5 µs
7.5 µs
10 µs
[U4]
[U4][U4]
[U4=U2][U2=U1][U2][U2=U1]
[U2=U1][U2=U1][U2][U2]
MD13|R4 5′-UUUU-3′
[ ][U2=U3]
[ ]
[U3][U3][U2/U3][U2/U3][U2]
0 µs
2.5 µs
5 µs
7.5 µs
10 µs
[U2][U2]
[ ][U2=U3][U2][U2=U3][ ]

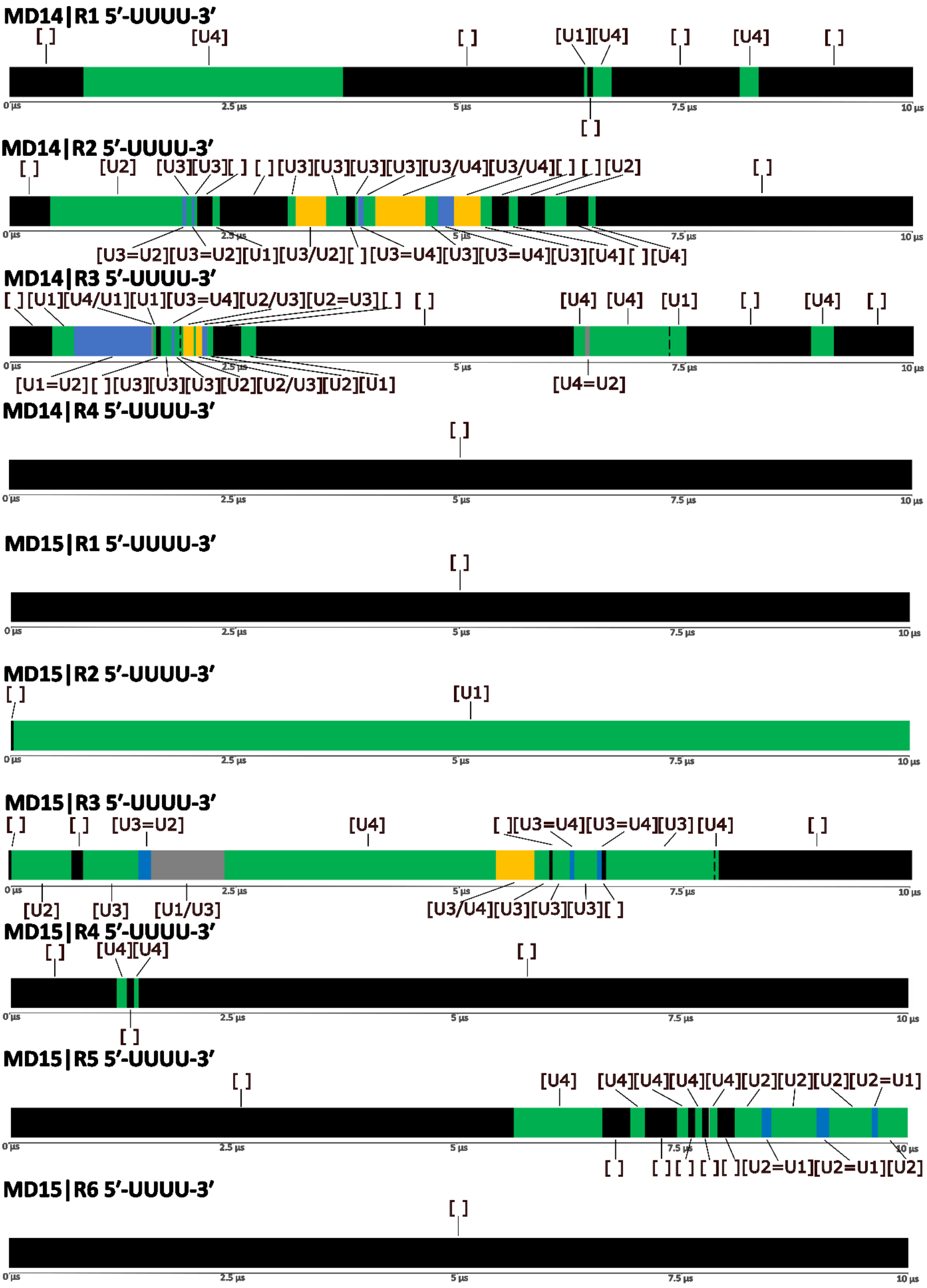


Figure S3: **Time development of the binding pocket state in all MD simulations of the TbRGG2 RRM complexes.** Each horizontal bar corresponds to a single MD simulation trajectory. The meaning of the different colors and of the labels above and below the graphs is explained in the legend at the top.

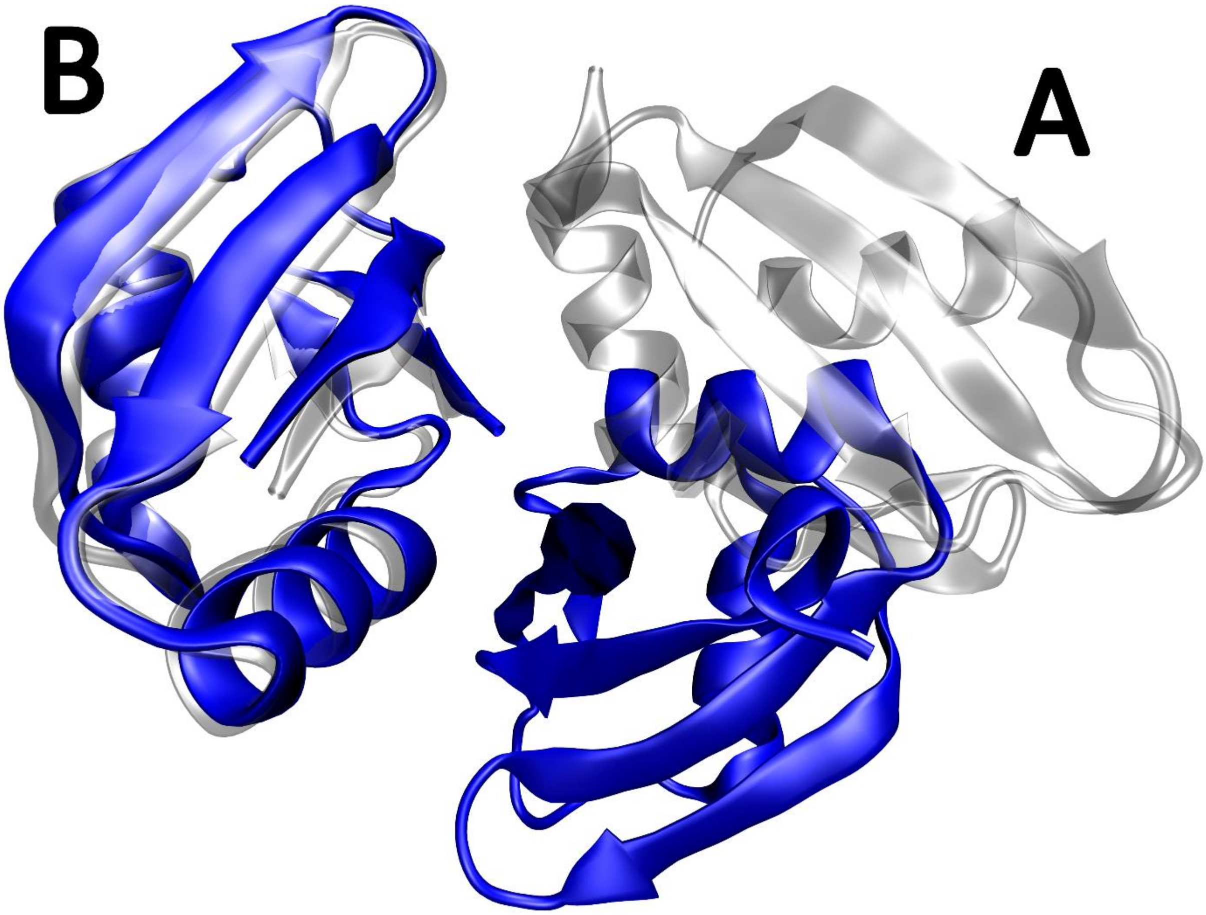


Figure S4: **MD simulations of the putative TbRGG2 RRM homodimer.** Superposition of the putative TbRGG2 RRM protein homodimer as shown in the experimental structure (chain A/B; grey) and after ~4µs of simulations (blue). Note that similar results were observed when simulating chains A/D.

### Supplementary Information Tables

Table S1. **List of supplementary MD simulations**.

| # | RNA sequence | Length (μs) |
|---|---|---|
| MDS1 | -[a] | 2 |
| MDS2 | -[b] | 2 |
| MDS3 | 5′-UUU-3′[a] | 2/2 |
| MDS4 | -[c] | 10/10 |

[a]Only chains A/B (putative homodimer formed by the RRMs) were simulated.
[b]Only chains A/D (putative homodimer formed by the RRMs) were simulated.
[c]Only chain B (RRM) was simulated.

Table S2: **Population of intermolecular H-bonds in the single uridine state of the binding pocket in selected MD simulations.** Only H-bonds with population higher than 5% are shown. The H-bonds are color-coded for clarity.

| **MD2\|R1** | | | | **MD4\|R1** | |
|---|---|---|---|---|---|
| **[U2]** (109 450 frames) | | **[U1]** (32 100 frames) | | **[U1]** (11 000 frames) | |
| **Hydrogen bond** | **Population** | **Hydrogen bond** | **Population** | **Hydrogen bond** | **Population** |
| Cys231(O) - U2(N3) | 68.17% | Cys231(O) - U1(N3) | 70.34% | Cys231(O) - U1(N3) | 76.22% |
| U2(O2) - Cys231(N) | 12.96% | U1(O2) - Cys231(N) | 48.41% | Thr229 (O) - U1(O2′) | 32.04% |
| U2(O4) - Lys232(NZ) | 8.54% | U1(O4) - Lys232(NZ) | 5.28% | U1(O2) - Cys231(N) | 30.66% |
| Thr229(O) - U2(O2′) | 6.17% | | | U1(O4) - Lys232(NZ) | 5.72% |

| **MD4\|R2** | | **MD4\|R4** | |
|---|---|---|---|
| **[U2]** (37 900 frames) | | **[U1]** (35 700 frames) | |
| **Hydrogen bond** | **Population** | **Hydrogen bond** | **Population** |
| Cys231(O) - U2(N3) | 84.24% | Cys231(O) - U1(N3) | 80.68% |
| U2(O2) - Cys231(N) | 33.25% | U1(O2) - Cys231(N) | 40.28% |
| Thr229 (O) - U2(O2′) | 31.78% | Thr229 (O) - U1(O2′) | 11.37% |
| U2(O4) - Lys232(NZ) | 6.14% | U1(O4) - Lys232(NZ) | 7.94% |

Table S3: **Population of intermolecular H-bonds in the vertical state of the binding pocket in selected MD simulations.** Only H-bonds with population higher than 5% are shown. The H-bonds are color-coded for clarity.

| **MD2\|R1** | | | | | |
|---|---|---|---|---|---|
| **[U2=U3]** (186 700 frames) | | **[U1=U2]** (106 900 frames) | | **[U0=U1]** (70 600 frames) | |
| **Hydrogen bond U3** | **Population** | **Hydrogen bond U2** | **Population** | **Hydrogen bond U1** | **Population** |
| Cys231(O) - U3(N3) | 66.80% | Cys231(O) - U2(N3) | 67.12% | Cys231(O) - U1(N3) | 66.49% |
| U3(O2) - Cys231(N) | 53.60% | U2(O2) - Cys231(N) | 56.56% | U1(O2) - Cys231(N) | 53.20% |
| U3(O4) - Lys232(NZ) | 5.85% | U2(O4) - Lys232(NZ) | 6.27% | U1(O4) - Lys232(NZ) | 5.46% |
| **Hydrogen bond U2** | **Population** | **Hydrogen bond U1** | **Population** | **Hydrogen bond U0** | **Population** |
| Cys231(O) - U2(N3) | 60.89% | Cys231(O) - U1(N3) | 57.98% | Cys231(O) - U0(N3) | 58.69% |
| U2(O4) - Val233(N) | 18.82% | U1(O4) - Lys232(NZ) | 8.67% | U0(O4) - Val233(N) | 8.67% |
| | | U1(O4) - Val233(N) | 6.71% | U0(O4) - Lys232(NZ) | 5.37% |
| **MD3\|R1** | | | | **MD4\|R1** | |

| **[U2=U3]** (29 800 frames) | | **[U1=U2]** (25 100 frames) | | **[U1=U2]** (97 500 frames) | |
|---|---|---|---|---|---|
| **Hydrogen bond U3** | **Population** | **Hydrogen bond U2** | **Population** | **Hydrogen bond U2** | **Population** |
| Cys231(O) - U3(N3) | 62.65% | Cys231(O) - U2(N3) | 59.57% | Cys231(O) - U2(N3) | 65.03% |
| U3(O2) - Cys231(N) | 46.43% | U2(O2) - Cys231(N) | 35.85% | U2(O2) - Cys231(N) | 54.65% |
| U3(O4) - Lys232(NZ) | 6.29% | U2(O4) - Lys232(NZ) | 7.35% | U2(O4) - Lys232(NZ) | 7.29% |
| **Hydrogen bond U2** | **Population** | **Hydrogen bond U1** | **Population** | **Hydrogen bond U1** | **Population** |
| Cys231(O) - U2(N3) | 61.95% | Cys231(O) - U1(N3) | 58.52% | Cys231(O) - U1(N3) | 55.78% |
| U2(O4) - Val233(N) | 8.63% | U1(O4) - Val233(N) | 9.43% | U1(O4) - Val233(N) | 8.69% |
| | | U1(O4) - Lys232(NZ) | 6.03% | U1(O4) - Lys232(NZ) | 5.19% |
| **MD4\|R1** | | **MD4\|R2** | | | |
| **[U0=U1]** (11 500 frames) | | **[U1=U2]** (8 100 frames) | | **[U2=U3]** (693 000 frames) | |
| **Hydrogen bond U1** | **Population** | **Hydrogen bond U2** | **Population** | **Hydrogen bond U3** | **Population** |
| U1(O2) - Cys231(N) | 60.81% | Cys231(O) - U2(N3) | 71.62% | Cys231(O) - U3(N3) | 66.65% |
| Cys231(O) - U1(N3) | 59.18% | U2(O2) - Cys231(N) | 46.90% | U3(O2) - Cys231(N) | 56.97% |
| Thr229(O) - U1(O2′) | 24.40% | Thr229(O) - U2(O2′) | 20.59% | U3(O4) - Lys232(NZ) | 7.10% |
| **Hydrogen bond U0** | **Population** | **Hydrogen bond U1** | **Population** | **Hydrogen bond U2** | **Population** |
| Cys231(O) - U0(N3) | 50.21% | U1(O4) - Lys232(NZ) | 19.01% | Cys231(O) - U2(N3) | 58.72% |
| U0(O4) - Lys232(NZ) | 17.24% | | | U2(O4) - Val233(N) | 6.68% |
| | | | | U2(O4) - Lys232(NZ) | 6.35% |
| **MD4\|R4** | | | | | |
| **[U1=U2]** (2 700 frames) | | **[U0=U1]** (161 600 frames) | | | |
| **Hydrogen bond U2** | **Population** | **Hydrogen bond U1** | **Population** | | |
| Cys231(O) - U2(N3) | 58.04% | Cys231(O) - U1(N3) | 67.92% | | |
| U2(O2) - Cys231(N) | 34.07% | U1(O2) - Cys231(N) | 63.08% | | |
| U2(O4) - Lys232(NZ) | 10.41% | U1(O4) - Lys232(NZ) | 5.95% | | |
| **Hydrogen bond U1** | **Population** | **Hydrogen bond U0** | **Population** | | |
| Cys231(O) - U1(N3) | 47.04% | Cys231(O) – U0(N3) | 57.67% | | |
| U1(O4) - Val233(N) | 18.22% | U0(O4) - Val233(N) | 8.22% | | |
| | | U0(O4) - Lys232(NZ) | 6.99% | | |

Table S4: **Population of intermolecular H-bonds in the horizontal state of the binding pocket in selected MD simulations.** Only H-bonds with population higher than 5% are shown. The H-bonds are color-coded for clarity.

| **MD3\|R3** | | **MD3\|R6** | | **MD4\|R5** | |
|---|---|---|---|---|---|
| **[U2/U3]** (33 400 frames) | | **[U2/U3]** (15 600 frames) | | **[U1/U2]** (28 400 frames) | |
| **Hydrogen bond U3** | **Population** | **Hydrogen bond U3** | **Population** | **Hydrogen bond U2** | **Population** |
| U3(O4) – Lys219(NZ) | 44.07% | U3(O4) – Lys219(NZ) | 39.59% | Thr229(O) – U2(O2′) | 64.06% |
| Val228(O) – U3(N3) | 18.39% | Val228(O) – U3(N3) | 27.66% | Val228(O) – U2(N3) | 40.65% |
| Thr229(O) – U3(O2′) | 11.19% | | | U2(O4) – Lys219(NZ) | 16.31% |
| **Hydrogen bond U2** | **Population** | **Hydrogen bond U2** | **Population** | **Hydrogen bond U1** | **Population** |
| Cys231(O) – U2(N3) | 72.74% | Cys231(O) – U2(N3) | 79.55% | Cys231(O) – U1(N3) | 70.49% |
| U2(O4) - Lys232(NZ) | 13.62% | U2(O4) - Lys232(NZ) | 24.03% | U1(O4) - Lys232(NZ) | 9.79% |

### Supplementary Information References


1. Krepl, M., Pokorná, P., Mlýnský, V., Stadlbauer, P. and Šponer, J. (2022) Spontaneous Binding of Single-stranded RNAs to RRM Proteins Visualized by Unbiased Atomistic Simulations with a Rescaled RNA Force Field. *Nucleic Acids Res.*, **50**, 12480–12496.
2. Travis, B., Shaw, P.L.R., Liu, B., Ravindra, K., Iliff, H., Al-Hashimi, Hashim M. and Schumacher, M.A. (2018) The RRM of the kRNA-editing Protein TbRGG2 Uses Multiple Surfaces to Bind and Remodel RNA. *Nucleic Acids Res.*, **47**, 2130-2142.
3. Eisenberg, D., Schwarz, E., Komaromy, M. and Wall, R. (1984) Analysis of membrane and surface protein sequences with the hydrophobic moment plot. *J. Mol. Biol.*, **179**, 125-142.
